# A dramatic protein fold switch powers a bactericidal nanomachine

**DOI:** 10.64898/2026.01.28.702463

**Authors:** Yao He, Annie Si Cong Li, Xiaoying Cai, Shoichi Tachiyama, Rajeev Kumar, Devlina Chakravarty, Lauren L. Porter, Jun Liu, Alan R. Davidson, Z. Hong Zhou

**Affiliations:** Department of Microbiology, Immunology and Molecular Genetics, University of California, Los Angeles (UCLA), Los Angeles, CA, USA; The California NanoSystems Institute (CNSI), University of California, Los Angeles (UCLA), Los Angeles, CA, USA; Department of Biochemistry, Department of Molecular Genetics, University of Toronto, Toronto, Ontario, Canada; Microbial Sciences Institute, Yale University, West Haven, CT 06516, USA; Department of Microbial Pathogenesis, Yale School of Medicine, New Haven, CT 06536, USA; National Library of Medicine, National Institutes of Health, Bethesda MD 20894, USA; National Heart Lung and Blood Institute, National Institutes of Health, Bethesda MD, 20894, USA

## Abstract

Fold switching, where a protein region interconverts between entirely distinct three-dimensional structures, is emerging as vital for certain protein functions. Here, we report a remarkable example in the F7 pyocin, a phage tail-like bactericidal nanomachine. Cryogenic electron microscopy and tomography reveal that a 163-residue segment of the central tail fiber undergoes a dramatic transition—from a trimeric α-helical coiled-coil to a triangular β-prism—upon binding to the bacterial cell surface. This massive fold switch remodels the tail tip, ejects the internal tape measure protein, and drives membrane puncture. Site-directed mutations that selectively destabilize the β-prism conformation completely abolish bactericidal activity without impairing particle assembly, implying that the energy released during this transition powers penetration. AlphaFold-based analyses further predict similar large-scale coiled-coil to β-prism switches in diverse non-contractile phage tails. This discovery reveals a sophisticated, ATP-independent strategy for microbial warfare and opens exciting possibilities for engineering next-generation bacteriocins to combat multidrug-resistant pathogens.

## Main

Folded proteins are generally expected to adopt a single unique three-dimensional structure under physiological conditions. However, in recent years, proteins with domains (50 to 75 residues) that markedly change secondary structure as part of their normal function have been described^1,2^. The role and importance of these fold-switching phenomena have been characterized in just a few cases^3,4^. Here, we describe a dramatic fold switch that powers the bactericidal action of the F-type pyocin, a bacteriophage (phage) tail-like nanomachine produced by *Pseudomonas aeruginosa* (*Pae*).

As part of their normal life cycle, strains of *Pae* produce two types of phage tail-like bactericidal nanomachines, known as R-type and F-type pyocins, which selectively kill other strains of the same bacterial species^5^. These complexes have evolved from phages, viruses that infect bacteria, retaining their tail structures while losing the viral capsid and genome^5–7^. R-type pyocins are related to contractile tails, whereas F-type are related to non-contractile tails^5–8^. The study of pyocins is of great interest because their potent antibacterial activity may be exploited for the development of novel therapeutics against antibiotic-resistant bacteria^5^. In both contractile phages and R-type pyocins, the force provided by tail contraction drives the tail tube through the target cell envelope^6,7,9^. By contrast, the mechanism by which F-type pyocins and other non-contractile tails, which are found in nearly 60% of all known phages^10^, puncture the target cell envelope remains poorly understood. These tails lack an obvious means to generate the energy required for this process.

In this study, we have determined the first atomic structures of an F-type pyocin in two functional states. Upon binding cognate receptors, the central tail fiber undergoes a dramatic fold switch, with a 163-residue coiled-coil transitioning into a β-prism structure. This fold switch is accompanied by remodeling of the tail tip and ejection of the tape measure protein, which is enclosed within the tail tube. Mutations that sterically inhibit this transition result in F-type pyocins that assemble normally but lack bactericidal activity, suggesting that the stored conformational energy released during the fold switching process drives membrane puncture. AlphaFold^11^ predictions suggest that this mechanism is conserved among diverse non-contractile phage tails.

### Overall structure of the F7 pyocin tube and tail tip

The F-type pyocins of *Pae* are classified into 11 groups, which differ only in the sequences of their proteins involved in determining strain bactericidal specificity^8^. To determine how these entities assemble and engage target cells, we used single-particle cryogenic electron microscopy (cryo-EM) to determine the high-resolution structure of a member from the F7 pyocin group. The acquired micrographs yielded four density maps (Extended Data Fig. 1) that were computationally montaged to create a complete structure of the F7 pyocin (Fig. 1a–d; Supplementary Video 1). This structure is similar to other non-contractile phage tail structures^12–14^, consisting of a long tube capped at the distal end by the tail terminator protein (TrP) (Fig. 1b,e). The proximal end of the tube is attached to the tail tip complex, which contains five proteins: the distal tail (DTP), the tape measure (TMP), the tail hub (THP), the tail hub internal (THIP), and the central fiber (CFP) (Fig. 1c,f). Finally, three trimeric side fibers (SFP) interact with a coiled-coil trimeric portion of the central fiber (Fig. 1d,g). The structures of the tail terminator, tube, tip, and fiber regions of the F7 pyocin were determined at resolutions of 3.1 Å, 2.9 Å, 2.9 Å, and 3.1 Å, respectively, after applying the appropriate symmetries during reconstructions (Extended Data Figs. 1 and 2; Supplementary Table 1). All proteins encoded in the F-type pyocin genomic region (Fig. 1a) were located within the F7 pyocin structure except for PyoF3 and PyoF4, the tail assembly chaperones that are not found in phage tails, and PyoF8, a protein required for phage tail assembly but never detected as a tail component. PyoF11 and PyoF12 were also not identified as will be discussed below.

**Fig. 1.**
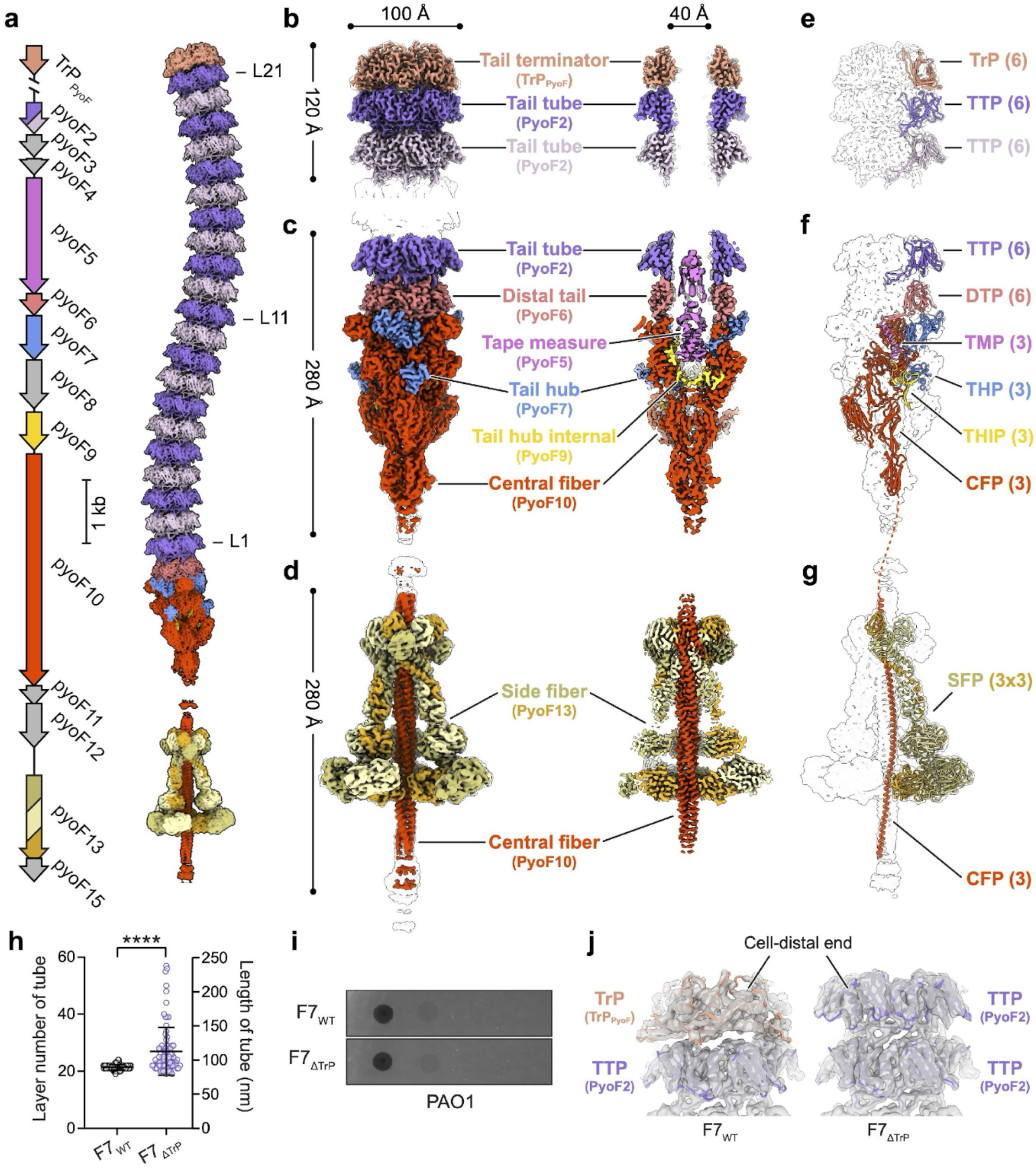
| Overall structure of F7 pyocin in the pre-ejection state. **a,** Organization of F7 pyocin genes. Genes colored in gray correspond to proteins that are not resolved in the cryo-EM reconstructions. The composite cryo-EM density map of the F7 pyocin in the pre-ejection state is shown on the right. Structural subunits are colored according to the gene map. **b–d,** Cryo-EM maps of the tail terminator (**b**), tip (**c**), and fiber (**d**) complexes of F7 pyocin, with corresponding sectional views shown on the right. **e–g,** Atomic models of the tail terminator (**e**), tip (**f**), and fiber (**g**) complexes of F7 pyocin, with protein assignments and corresponding copy numbers indicated. **h,** Ten-fold dilutions of F7_WT_ and F7_ΔTrP_ lysates were spotted on a lawn of *P. aeruginosa* (*Pae*) strain PAO1. Bactericidal activity is indicated by zones of clearing. **i,** Statistics of the layer number and length of tail tube in F7_WT_ and F7_ΔTrP_ particles. Data are represented as mean ± STD. The sample sizes are 82 and 86 for F7_WT_ and F7_ΔTrP_ particles from three independent replicate EM grids, respectively. ****P < 0.0001 by unpaired *t*-test. **j,** Cryo-EM density maps (semi-transparent densities) of the cell-distal ends of F7_WT_ and F7_ΔTrP_ particles, encasing the corresponding atomic models (ribbons). Both maps were low-pass filtered to 5 Å for comparison.

**Fig. 2.**
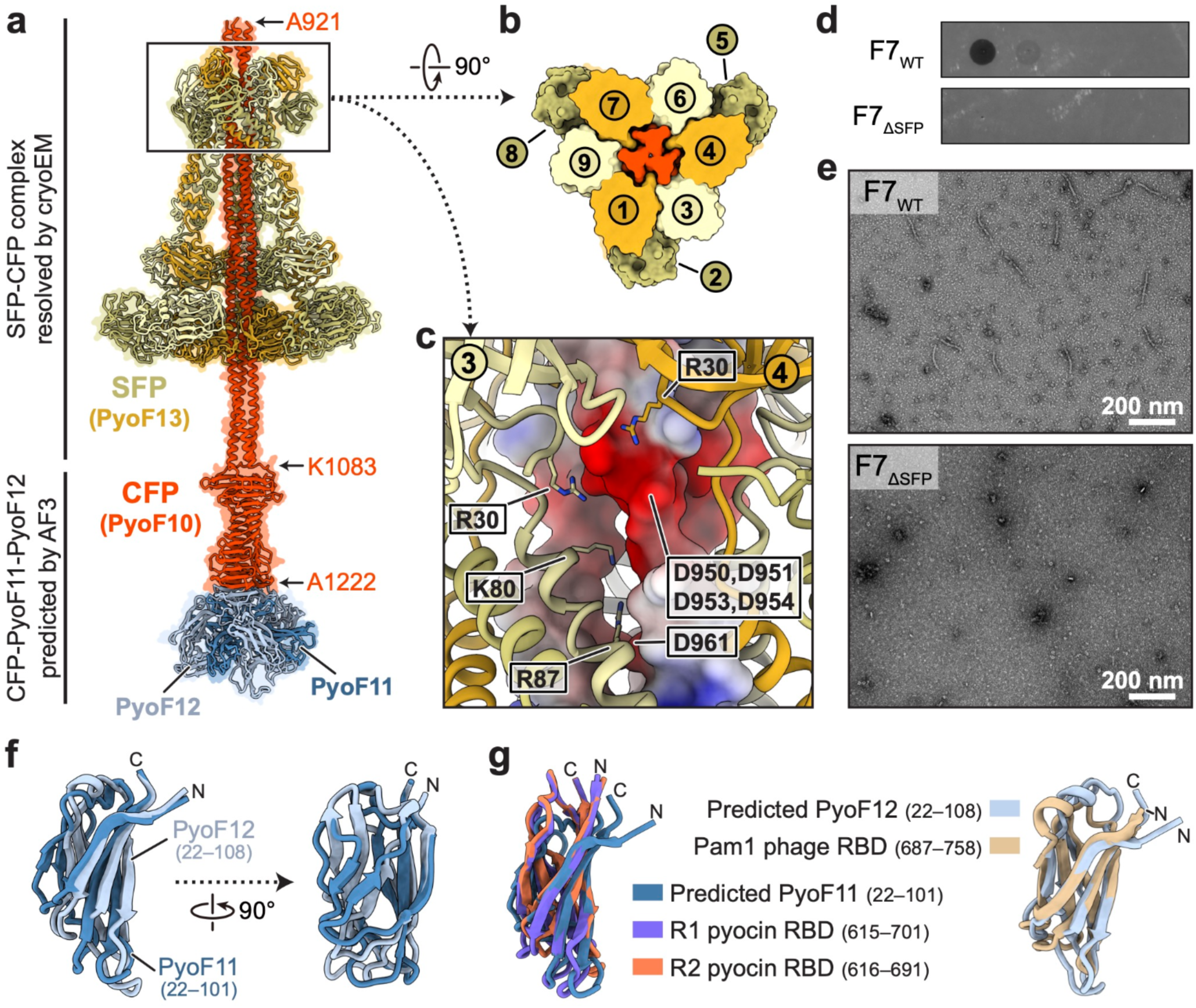
| Interactions between the central and side fiber proteins. **a,** Ribbon diagram of the fiber complex. The SFP-CFP complex was resolved by cryo-EM, and the C-terminal regions of the CFP in complex with PyoF11 and PyoF12 was modeled using AlphaFold3 (AF3) ^27^. **b**, Sectional view of the interface between the CFP and SFP, with nine SFP molecules numbered as indicated. **c,** Close-up view of the interface between the SFP and CFP. Positively charged residues of the SFP on the interface are shown as sticks. The CFP is shown as an electrostatic surface with negative charges in red, positive charges in blue, and neutral regions in white. **d,** Ten-fold dilutions of F7_WT_ and F7_ΔSFP_ lysates were spotted on a lawn of *Pae* strain PAO1. **e,** Representative negative-stain EM images of F7_WT_ and F7_ΔSFP_ lysates. **f,** Structural alignments of the potential cell surface receptor binding domains (RBDs) of PyoF11 and PyoF12. **g,** Comparison of the RBDs of PyoF11 with those of R-type pyocins (R1 pyocin RBD PDB: 6CL5; R2 pyocin RBD PDB: 6CL6) ^76^, and the RBDs of PyoF12 with that of phage Pam1 (PDB: 7EEA) ^77^.

In cryo-EM micrographs, the tail tube of the F7 pyocin appears as a long and flexible structure (Extended Data Fig. 1a); this flexibility is reflected in the density map obtained through asymmetric reconstruction (Extended Data Figs. 1b and 3a) and further supported by 3D variability analysis^15^ (3DVA) (Supplementary Video 2). The tail tube length is highly consistent among F7 pyocin particles, averaging 84 nm with 21 layers of hexameric tail tube protein (TTP) rings (Fig. 1a,h). Like the N-terminal domain of the phage lambda TTP^13^, the F7 pyocin TTP features a bent β-sandwich domain flanked by an α-helix (Extended Data Fig. 3b). This “tail tube fold” is conserved in the tail tube proteins of all long-tailed phages^16^.

**Fig. 3.**
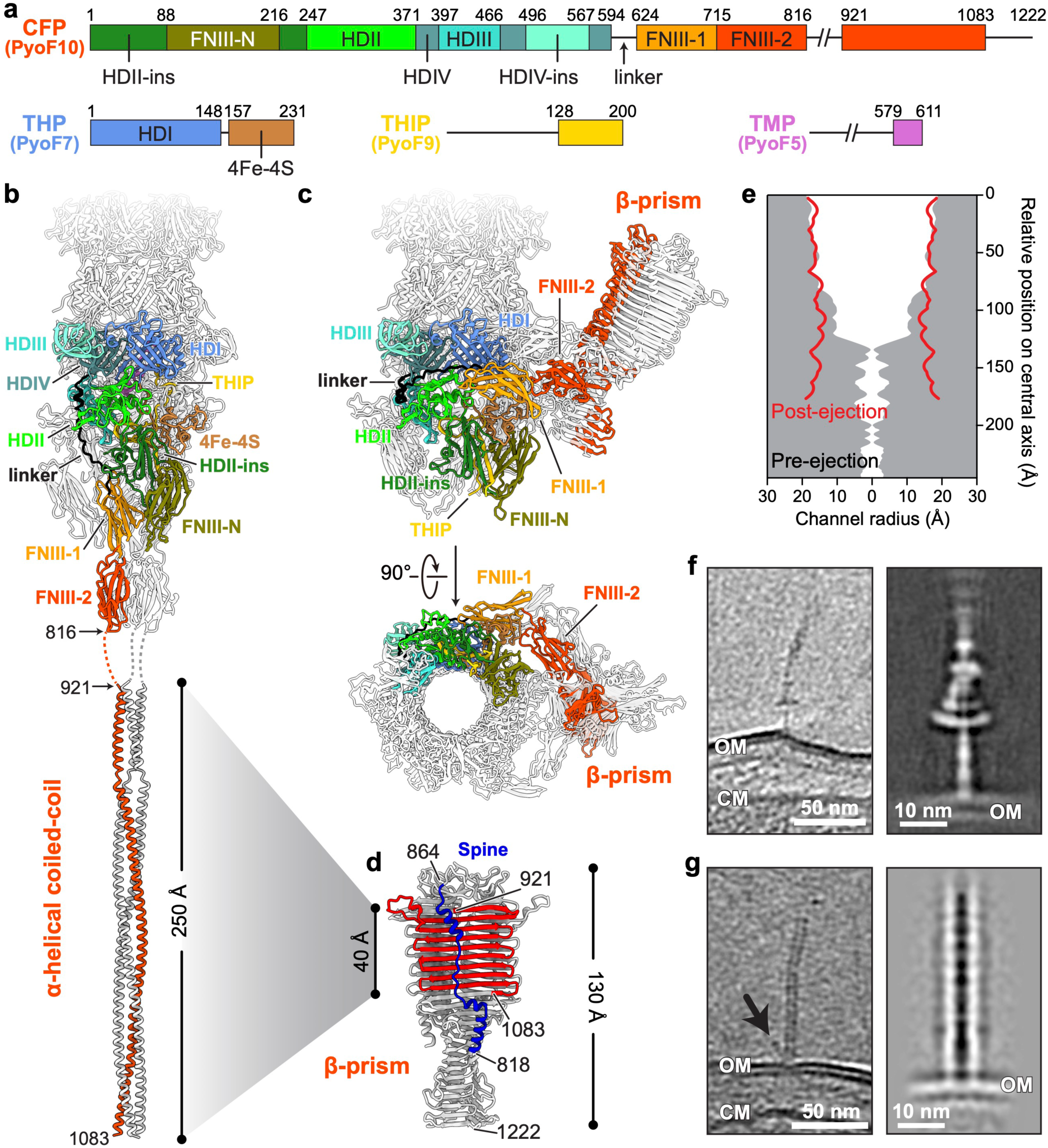
| The central fiber undergoes a massive fold switching. **a,** Linear schematic of the domain organizations of the CFP, THP, THIP, and TMP. **b–c,** Ribbon diagrams of the tail tip and central fiber in the pre-ejection (**b**) and post-ejection (**c**) states. Individual domains are colored as in **a**. The C-terminal coiled-coil of the CFP trimer switches from α-helical coil-coil to a β-prism. **d,** Triangular β-prism of CFP trimer in the post-ejection state. Residues 921–1083, corresponding to the coiled-coil region in the pre-ejection state (**b**), are colored in red. Residues 818–864, which form a “spine” on the solvent-exposed surface of the β-prism, are colored in blue. **e,** Comparison of the lumen radii of tail tips in the pre- (white-grey boundaries) and post-ejection (red lines) states. The radii were calculated using HOLE^78^. For the pre-ejection state, TMPs were removed from the lumen before calculation. **f,** A slice image of a cryo-ET reconstruction of F7 pyocin in the pre-ejection state bound to *Pae* cell surface (left), and the subtomogram-averaged density map of the fiber complex (right). **g,** A slice image of a cryo-ET reconstruction of F7 pyocin in the post-ejection state on the *Pae* cell surface (left). The density corresponding to the CFP β-prism is indicated by an arrow. A subtomogram-averaged density map of F7 pyocin in the post-ejection state is shown on the right, highlighting the absence of the TMP. OM, outer membrane. CM, cell membrane.

The hexameric ring of the DTP acts as the interface between the tail tip and the proximal end of the tail tube (Fig. 1c,f). Despite a completely different sequence from the TTP, the DTP also adopts a tail tube fold as it does in other siphophages^17^ (Extended Data Fig. 3b). The bulk of the tail tip is comprised of the CFP, which displays a complex arrangement of multiple domains, and the THP (Extended Data Fig. 3c, d). Tail tube fold-like domains formed by the CFP and the THP interlace to form a ring with pseudo-hexameric symmetry that serves as an adaptor to link the 6-fold symmetry of the tail tube to the 3-fold symmetry of the tail tip (Extended Data Fig. 3e–h). A conserved C-terminal domain of the THP containing an iron-sulfur cluster likely stabilizes the tail tip through interactions with several CFP domains (Extended Data Fig. 3c, d). The THIP forms a meandering structure that extends through the interior of the tail tip (Fig. 1c,f; Extended Data Fig. 3i–l). Finally, the 33 C-terminal residues of the TMP form a trimeric helical arrangement inside tail tip, interacting primarily with the THIP (Fig. 1c,f; Supplementary Video 3).

### The F7 pyocin TrP is encoded at a distinct locus

In our previous work, we found that the F-type pyocin genomic region lacks a gene encoding a TrP^8^, which is present in all long-tailed phages^18^. Thus, it was surprising that the cryo-EM reconstruction of F7 pyocin revealed a hexameric protein ring, distinct from the TTP ring, capping the distal end of the tail tube (Fig. 1b,e). To identify this unknown protein, we used ModelAngelo^19^ to build an initial model into the density without providing sequence information. The resulting model was structurally similar to TrPs from diverse contractile and non-contractile phages (Extended Data Fig. 4a–f), and the predicted protein sequence shared 97% pairwise identity with the product of *PA0910* from the reference *Pae* strain PAO1. This protein, which is detectably similar in sequence to some phage TrPs (Extended Data Fig. 4g), is conserved across nearly all *Pae* strain genomes^20^. However, its gene is located approximately 290 kbp from the F-type pyocin gene operon. To confirm that it encodes the TrP, we constructed an in-frame deletion of the homologue of *PA0910* in our F7 pyocin production strain.

Although F7 pyocins produced by this mutant strain (F7_ΔTrP_) retained bactericidal activity comparable to that of wild type (F7_WT_) (Fig. 1h), the average length of F7_ΔTrP_ particles increased and the tail lengths showed greater variation (Fig. 1h; Extended Data Fig. 4h), as has also been observed for phage lambda mutants lacking the TrP^21,22^. Cryo-EM reconstruction further confirmed that F7 pyocin particles produced from the F7_ΔTrP_ strain lacked the TrP at their distal end (Fig. 1j). The phenomenon of a component of a phage-related bacterial entity being encoded in a separate genome location has been previously observed^23,24^, but it is uncommon.

### The F7 pyocin fibers are arranged in a unique manner

The C-terminal region of the CFP and the SFP have been strongly implicated in determining the bactericidal strain specificity of F-type pyocins^8^. Similar to structures of non-contractile phage tails^13,25^, the CFP (1222 residues) comprises the bulk of the F7 pyocin tail tip. The F7 pyocin CFP is unique compared to solved structures of siphophage CFPs in possessing a long trimeric coiled-coil structure formed by residues 921 to 1083 that extends distally from the tail tip (Fig. 2a). Another unique feature of the F7 pyocin tail fiber is the attachment of three sets of trimeric side fibers to a short segment of the coiled-coil region (Fig. 2a,b; Supplementary Video 3). Positively charged residues in the N-terminal domain of the side fibers (Arg30, Lys80, and Arg87) engage with a negatively charged gap within the coiled-coil structure of the CFP at residues Asp950, Asp951, Asp953, Asp954, and Asp961 (Fig. 2c). These electrostatic interactions may stabilize the SFP-CFP complex. Lysates of a *Pae* mutant bearing an in-frame deletion in the gene encoding the SFP (F7_ΔSFP_) displayed no bactericidal activity (Fig. 2d) and no intact F7 pyocin particles were observed by negative-stain EM (Fig. 2e). These results suggest that the CFP cannot maintain its coiled-coil architecture without stabilization through interaction with SFP trimers.

The 139 residues at the C-terminal end of the CFP were not resolved in our structure, nor were two additional proteins, PyoF11 and PyoF12. These proteins, which are encoded immediately downstream of the CFP, are essential for bactericidal activity and have been detected in F7 pyocin particles through mass spectrometry analysis^8,26^. Since these proteins and the C-terminal region of the CFP have been implicated in strain bactericidal specificity, we used AlphaFold3^27^ (AF3) to investigate whether PyoF11 and PyoF12 may form a complex with the unresolved, C-terminal end of the CFP. AF3 predicts that these two proteins attach to the C-terminus of the CFP (Fig. 2a) and adopt structures that are similar to the C-terminal receptor binding domains (RBDs) of the fibers of R-pyocin and phage Pam1 (Fig 2f,g). In models of the CFP:PyoF11:PyoF12 complexes predicted for other F-type pyocin groups, PyoF11 and PyoF12 are also seen to bind to the end of the CFP and form similar structures to the F7 pyocin proteins (Extended Data Fig. 5a) even though these homologues display low sequence identity (< 25%) (Extended Data Fig. 5b). Predictions of phage encoded PyoF11 and PyoF12 homologues in complex with respective CFPs also yielded similar complexes (Extended Data Fig. 5a). Despite relatively low confidence values among these AF3 generated models, the similarity among structures of diverse homologues supports the conclusion that PyoF11 and PyoF12 serve as cell surface binding domains attached to the C-terminal end of the CFP.

### The central fiber undergoes a large fold switch

Among the F7 pyocin particles observed in the cryo-EM micrographs, approximately half displayed a dramatically different structure from that described above. Symmetry relaxation from C3 to C1 during cryo-EM reconstruction of these particles resolved an asymmetric density map of a second tail tip structure at 3.6 Å resolution (Fig. 3a,c; Extended Data Figs. 1b and 6a,b). The most remarkable feature of this structure is that the whole 163-residue α-helical coiled-coil region of the CFP is converted to a β-prism structure comprised of three large anti-parallel β-sheets (Fig. 3b–d). Residues 818–921 of the CFP, which were not resolved in our structure described above, form a “spine” structure composed of extended and helical regions on the outside surface of the β-prism (Fig. 3d). The β-prism is positioned to one side of the tail tube so that the bottom of the tail tube becomes opened, allowing ejection of the TMP (Fig. 3c,e). TMP ejection is evidenced by the loss of internal density from the tail tube (Extended Data Fig. 6a). This second structural state of the F7 pyocin will be referred to as the post-ejection state, while the state described above will be referred to as the pre-ejection state.

In the post-ejection state, the formation of the β-prism and its displacement to the side of the tail tip is accompanied by a complicated series of conformational changes in domains of the CFP. The CFP is made up of multiple structurally conserved domains that have, by convention, been called Hub Domains (HDII to HDIV) ^13,25^ (Fig. 3a). The HDII, HDIII, and HDII-ins domains of the CFP along with the iron-sulfur cluster domain of the THP undergo a rigid rotation and move outward toward the tail tip axis (Fig. 3b,c; Supplementary Video 4), leading to a diameter expansion of the tail tip interior, from approximately 25 Å to 38 Å (Fig. 3e). The FNIII-N domain is repositioned between neighboring HDII-ins domains, stabilizing the open-tip conformation (Fig. 3c). The FNIII-1 and FNIII-2 domains are lifted, removing the closure at the tail tip (Fig. 3c; Extended Data Fig. 6a, c). The lifted FNIII-1 and FNIII-2 trimer dissociates into three subunits: two relocate to one side of the tail tip, while the third shifts to the opposite side (Extended Data Fig. 6c). This reconfiguration of these FNIII domains is facilitated by a flexible linker connecting HDIV to the FNIII-1 domain (Fig. 3b,c; Extended Data Fig. 6c).

Conformational change of the THIP leads to the opening of the tail tip through removal of the trimeric plug formed by this protein inside the tail tip (Fig. 3c; Extended Data Fig. 6d; Supplementary Video 4). These structural rearrangements are localized to the N-terminus of THIP, where residues 171 and earlier undergo a downward movement (Extended Data Fig. 6d). Although the THIP is traceable only from residue 158 onward, AF3^27^ predicts the presence of two hydrophobic α-helices within residues 94–142 (Extended Data Fig. 6e). These helices are also predicted as transmembrane helices by TMHMM 2.0^28^, suggesting their possible insertion into the outer membrane of the target bacterium (Extended Data Fig. 6e), similar to the THIP of phage lambda (gpI) ^29^.

The SFPs and PyoF11/F12 were not resolved in the post-ejection state, possibly due to their detachment from the tail tip during the conformational change. However, C-terminal residues of the CFP (residues 1084–1222) that could not be resolved in the pre-ejection state were clearly identified in the post-ejection state (Fig. 3d). This region forms the bottom of the β-prism structure (residues 1083–1119), followed by a short intertwined β-helix (residues 1119–1167), and then another narrower β-prism structure encompassing the rest of the protein (Fig. 3d; Supplementary Video 5).

### Cryo-ET shows the pre- and post-ejection states *in situ*

To determine whether the pre- and post-ejection states observed in our single particle cryo-EM study actually form during the interaction of F7 pyocins with cells, we performed cryogenic electron tomography (cryo-ET) of F7 pyocin particles mixed with *Pae* cells (Fig. 3f; Extended Data Fig. 7a). Subtomogram averaging of the region interacting with the bacterial surface revealed a structure very similar to the tail fiber in the pre-ejection state: a cone-shaped density corresponding well to the structure of the side fibers traversed by a rod-like density corresponding to the coiled-coil region of central fiber (Extended Data Fig. 7b, c; Supplementary Table 2). Densities likely corresponding to PyoF11 and PyoF12 attached to the C-terminus of the CFP were observed (Extended Data Fig. 7c). The structure at the end of the coiled-coil appears to interact with the surface of the *Pae* outer membrane (Extended Data Fig. 7b, c). In these density slices of the reconstructed tomograms, the bottom of the tail tube lies ∼30 nm away from the outer membrane (Fig. 3f; Extended Data Fig. 7a).

Cryo-ET also revealed structures similar to the post-ejection state described above that interact with *Pae* cell surface (Fig. 3g; Extended Data Fig. 7d). A structure resembling the β-prism region of the CFP was seen to the side of the tube, which is approximately 10 nm in length, comparable to the ∼13 nm observed in our single-particle cryo-EM reconstructions (Fig. 3e). The number of post-ejection particles in the reconstructed tomograms was insufficient to permit subtomogram averaging of the prism structure; however, subtomogram averaging of the tube region clearly revealed the absence of TMP density (Fig. 3g; Extended Data Fig. 7e; Supplementary Table 2). These data strongly support the biological relevance of the post-ejection state observed in our single-particle structure cryo-EM studies, and of fold switching process that is required to form this state.

### Blocking the α-helix-to-β-prism transition abrogates bactericidal activity

To assess the functional importance of the coiled-coil to β-prism transition, we sought to introduce amino acid substitutions into the CFP that would decrease the chance of its occurrence. We reasoned that substitutions reducing the stability of the β-prism without affecting the stability of the coiled-coil would energetically favor the pre-ejection state and discourage fold switching. For this purpose, we identified more than 20 positions that are very tightly packed in the corners of the β-prism structure where the sheets formed by the individual CFP subunits interact (Fig. 4a). These positions are occupied exclusively by small amino acids, mostly Ala, but also some Val, Gly and Thr, likely because the tight packing at these prism corners does not provide space for larger residues (Fig. 4b). These residue positions are fully exposed in the coiled-coil structure; thus, substituting them should only affect stability of the β-prism. We introduced mutations that simultaneously replaced four of these residues (Ala987, Ala995, Val1010, and Ala1017; Extended Data Fig. 8a) with Gln (F7_CFP-4Q_). We expected that the greater polarity and surface area of the Gln side chain compared to Ala or Val^30,31^ would be destabilizing within the context of the tight interior packing of these positions in the β-prism. The stability of the coiled-coil should not be affected by these substitutions as these residues are exposed (Extended Data Fig. 8a) and Gln has a high propensity for helical conformation^32^. Lysates containing the wild-type or mutant (F7_CFP-4Q_) were spotted on a sensitive bacterial lawn. A zone of bacterial growth inhibition was observed for the wild-type lysate, but not the mutant lysate, showing that the mutant lacked bactericidal activity (Fig. 4c). Testing additional mutants showed that a A987Q/A995Q double mutant (F7_CFP-2Q_) (Fig. 4b) also lacked bactericidal activity (Fig. 4c). To demonstrate the specificity of this effect, we simultaneously substituted nearby residues (A998Q and A1011Q) that are exposed on the surface of both the β-prism and coiled-coil structures (Fig. 4b). This double mutant (F7_CFP-2QC_) displayed no reduction in bactericidal activity (Fig. 4c).

**Fig. 4.**
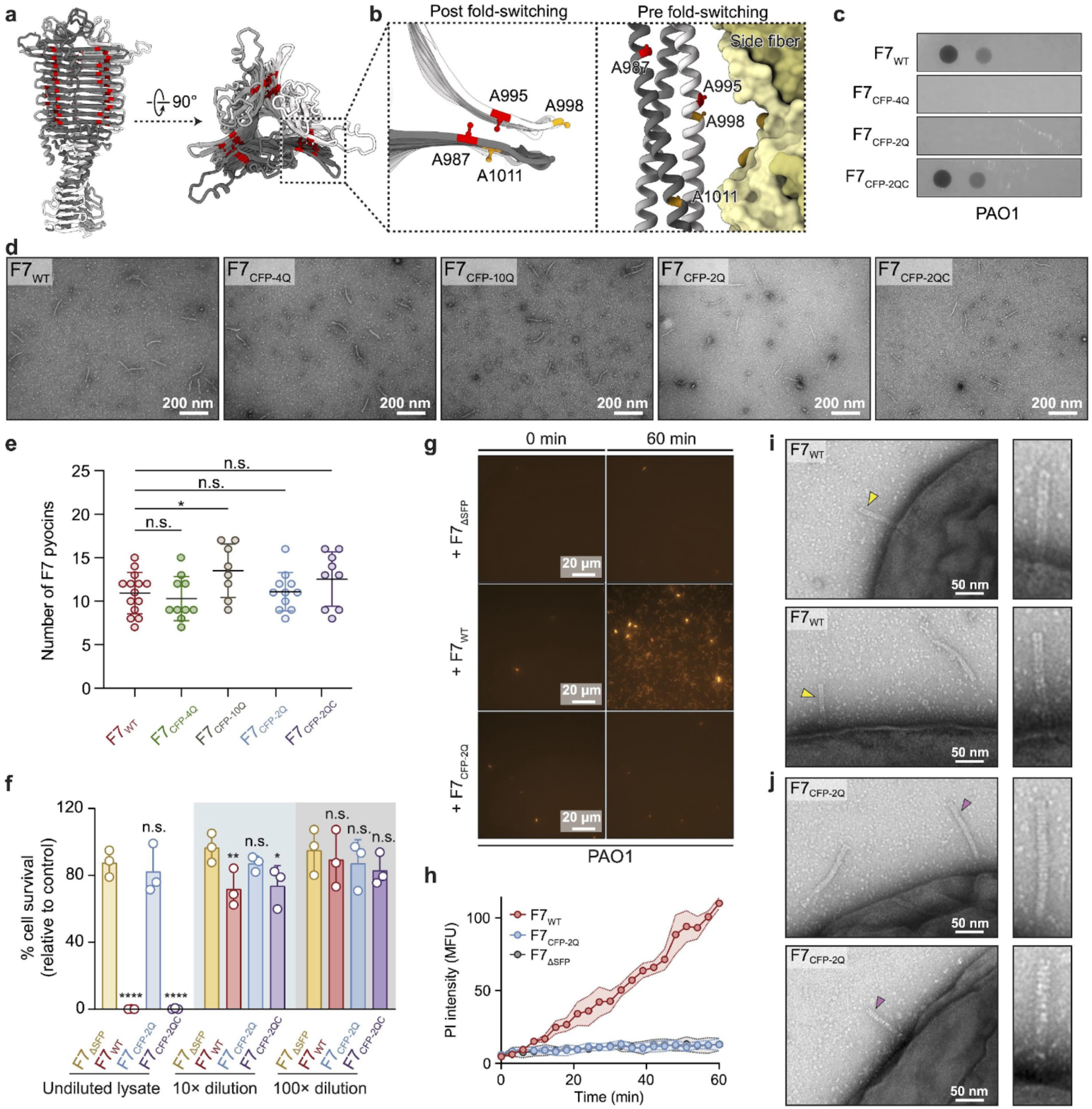
| Amino acid substitutions abrogate F7 pyocin bactericidal activity without affecting assembly. **a,** Ribbon diagram of the CFP β-prism. The locations of the small residues in the corners of the β-prism (typically Ala) are highlighted in red. **b,** Positions of residues substituted in the F7_CFP-2Q_ mutant (A987 and A995 in red) and F7_CFP-2QC_ mutant (A998 and A1011 in orange) in the β-prism structure (left) and coiled-coil (right) structure. **c,** Ten-fold dilutions of WT and mutant F7 pyocin lysates were spotted on a lawn of *Pae* strain PAO1. **d,** Representative negative-stain EM images of WT and mutant F7 pyocin lysates. **e,** Quantification of the number of assembled tails observed in multiple electron micrographs of WT and F7 pyocin mutants taken at random locations across carbon-coated grids. Statistical significance was measured using unpaired *t*-test against F7_WT_ as a control. \**p-*value = 0.04. n.s. = not significant, n=3. **f,** Quantitative cell killing of PAO1 cells by WT and mutant F7 pyocin lysates. Percent cell survival was calculated by comparing colony-forming units (CFUs) in pyocin-treated samples to those in untreated controls. The lysate of F7_ΔSFP_, which produces no assembled F7 pyocins, was tested for comparison. Asterisks show statistical significance relative to the corresponding F7_ΔSFP_ dilution, calculated by unpaired *t*-test. n.s. = not significant. *p-value = 0.026. ** p-value= 0.0035. ****p-value <0.0001. n=3. **g,** Time-lapse fluorescence microscopy images of PAO1 cells incubated with F7_WT_, F7_CFP-2Q_, or F7_ΔSFP_ in the presence of propidium iodide (PI). Cells were imaged over the course of 1 hour with a widefield epifluorescence microscope at 3 min intervals. Time points at 0 min and 60 min after adding F7 pyocins are shown. **h,** Quantification of the time-lapse fluorescence microscopy data. Mean PI fluorescence intensities (n=3) of cells treated with indicated lysate at different time points are shown. **i–j,** Representative negative-stain EM images of F7_WT_ (**i**) and F7_CFP-2Q_ mutant (**j**) particles interacting with PAO1 cells. Yellow arrows indicate “empty” F7 pyocins that are devoid of the internal electron density corresponding to the TMP. Purple arrows indicate “full” F7 pyocins with TMP. Zoom-in views of the arrow-marked particles are shown on the right.

We used negative-stain EM to verify that the abrogated bactericidal activities observed for the F7 pyocin mutants were not due to a defect in tail particle assembly. Examination of multiple randomly selected EM fields showed that lysates of both the F7_CFP-4Q_ and F7_CFP-2Q_ mutants contained comparable numbers of fully assembled tail particles to the lysate of F7_WT_ (Fig. 4d,e). Remarkably, even a lysate of a mutant with Gln substitutions at 10 β-prism corner positions (F7_CFP-10Q_) (Extended Data Fig. 8b) contained a similar number of tail-like particles as the F7_WT_ lysate (Fig. 4d,e). The tail tip complex in the pre-ejection state, characterized by its signature cone-shaped feature, was observed in particles from the lysates of all F7 mutants. These data clearly demonstrate that amino acid substitutions at key positions in the β-prism structure do not affect formation of the pre-ejection state.

Quantitative bactericidal assays were performed to confirm the effect of the F7_CFP-2Q_ mutant. In these assays, dilutions of F7 pyocin lysates were incubated with a defined number *Pae* cells, and surviving colony-forming units were counted. The F7_CFP-2Q_ mutant lysate elicited no detectable cell killing activity, similar to the lysate of F7_ΔSFP_ (Fig. 4f), which does not assemble into particles (Fig. 2e). By contrast, the F7_CFP-2QC_ mutant displayed the same level of activity as the F7_WT_ lysate (Fig. 4f). In summary, our mutant analysis supports the conclusion that these amino acid substitutions block the coiled-coil to β-prism switch, and that this switch is crucial for bactericidal activity.

### A β-prism destabilized mutant does not eject the TMP

During siphophage infection, the TMP is ejected from the tail tube and likely forms a channel through the periplasm and inner membrane to mediate DNA entry into the target bacterial cell^33–37^. For F-type pyocins, the TMP is similarly proposed to mediate puncturing of the inner membrane, leading to depolarization and cell death. Since the coiled-coil to β-prism switch appears to be required to open up the tail tip and allow TMP egress, we predicted that the F7_CFP-2Q_ mutant, which cannot undergo this transition, would be unable to mediate cell membrane puncturing. To directly detect cell membrane puncturing by the F7 pyocin, we added F7 pyocin lysates to *Pae* cells in the presence of propidium iodide (PI), a membrane-impermeable DNA intercalating dye that fluoresces upon DNA-binding^38^. Addition of F7_WT_ lysate resulted in a rapid increase in the level of PI fluorescence (Fig. 4g,h), indicating that pyocin-mediated membrane puncture allowed PI to enter the cells and bind to the bacterial nucleic acids. In contrast, adding the F7_CFP-2Q_ mutant lysate to cells under the same conditions led to no increase in fluorescence, demonstrating that destabilization of the β-prism conformation impairs the ability of the F7 pyocin to puncture the inner membrane. Similarly, lysate of the F7_ΔSFP_ mutant, which does form tail structures (Fig. 2e), also caused no increase in PI fluorescence (Fig. 4g,h). Together, these data directly show that amino acid substitutions designed to block the CFP fold-switching prevent F7 pyocin-mediated membrane puncturing.

To directly assess the effect of the F7_CFP-2Q_ mutant on TMP ejection, we incubated WT and mutant pyocins with *Pae* cells and examined their interactions by negative-stain EM. We found that 91% of cell-bound F7_WT_ pyocins displayed a loss of electron density in the lumen of the tail tube (Fig. 4i; Extended Data Fig. 8c), indicating TMP ejection. By contrast, all F7_CFP-2Q_ mutant pyocin particles that appeared to be in contact with cell surface, as judged by proximity and perpendicular orientation to the membrane, retained their TMP (Fig. 4j; Extended Data Fig. 8d). These data imply that inhibiting the switching from coiled-coil to β-prism prevents the conformational change in the tail tip required for TMP ejection and subsequent bactericidal activity.

### Coiled-coil to β-prism switching in other phage CFPs

We hypothesized that if fold switching plays a crucial role in the biological function of the F7 pyocin, then this phenomenon might be conserved in some non-contractile tailed phages. To address this issue, we performed a BLAST^39^ search using the final 400 residues of the F7 pyocin CFP as a query. From a database of 120 siphophages infecting Gram-negative bacteria, we identified 24 related but diverse CFPs (average pairwise identity of 29%, Supplementary Fig. 1). AF3^27^ predicted a triple α-helical coiled-coil for 70% of these regions and a β-prism for the other 30%. Remarkably, in four cases, a coiled-coil structure was predicted for one phage sequence, while a very similar sequence (50 to 80% identical) was predicted to fold into a β-prism (Extended Data Fig. 9a). All of these regions had very high coiled-coil propensity (probability > 0.95) as predicted by CoCoNat^40^ (Supplementary Table 3).

To systematically investigate the fold-switching nature of these CFPs, we used a newly developed implementation of AlphaFold2 (AF2), called CF-random^11^, that was designed to predict alternative protein conformations by sampling shallower multiple sequence alignments (MSAs). Combining predictions from both deep and shallow MSAs, CF-random predicted both β-prism (deep) and coiled-coil (shallow) conformations for 21 of 24 phage CFPs, and for the F7 pyocin CFP C-terminal region (Fig. 5a; Extended Data Fig. 9b; Supplementary Table 3). The predicted structures of the regions C-terminal to the fold-switching region are similar, with each of these non-switching regions commencing with three strands of β-prism (Fig. 5a; Extended Data Fig. 9b). This small region of β-prism structure may serve as the seed for switching of the coiled-coil region. The first 20–60 residues of the predicted coiled-coils of the phage CFPs do not switch to a β-prism structure but rather adopt a helical or extended “spine” structure that runs perpendicular to the strands comprising the β-prism (Fig. 5a; Extended Data Fig. 9b), similar to the “spine” observed in the cryo-EM structure of the F7 pyocin (Fig. 3d).

**Fig. 5.**
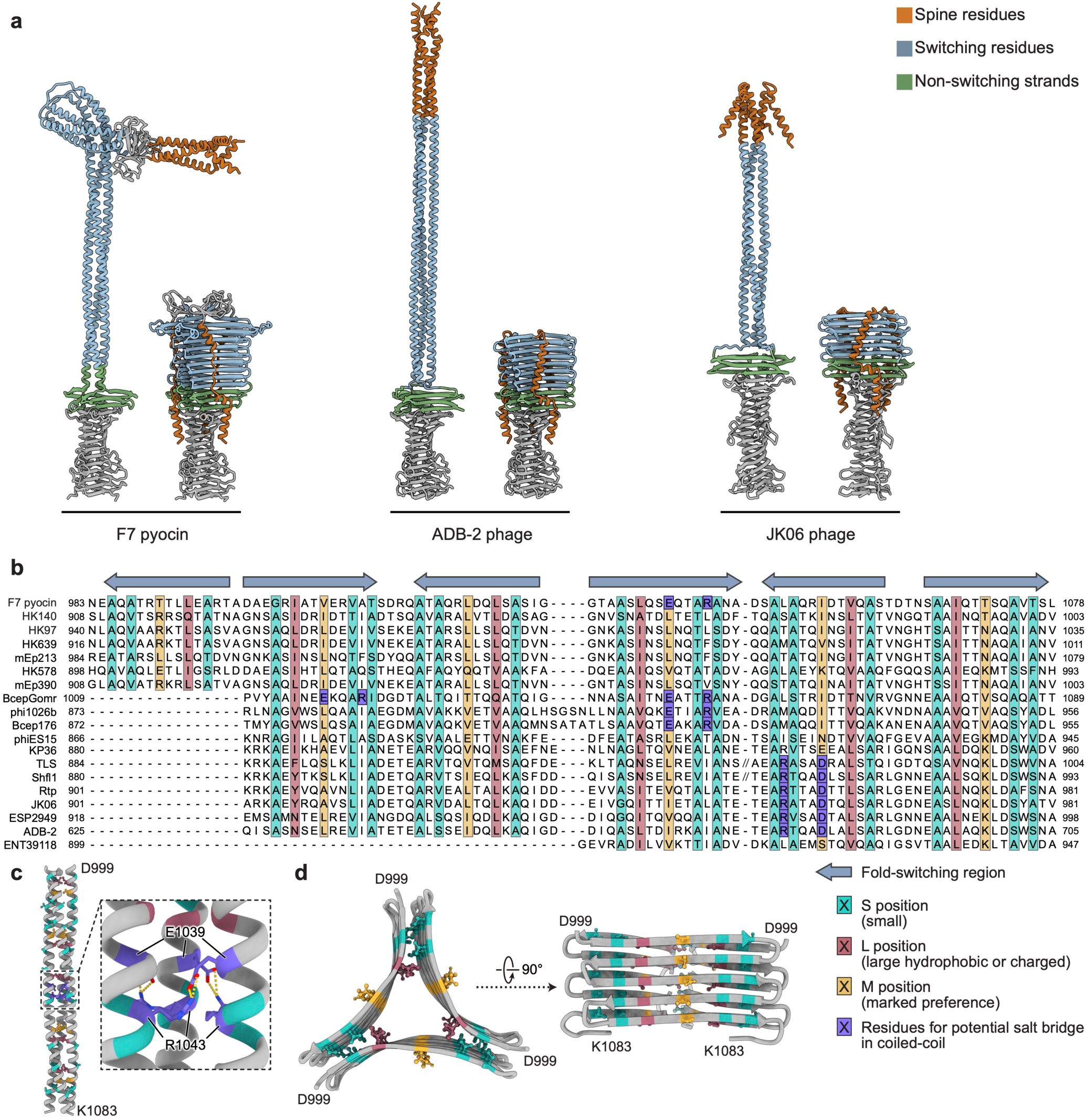
| Diverse phage central fiber proteins are predicted to undergo coiled-coil to β-prism fold switching. **a,** Fold-switched CFPs of the F7 pyocin and two phages predicted by CF-random^11^. Residues forming the “spine” are coloured in dark orange, the β-prism strands seen in the coiled-coil are coloured in blue, and non–fold-switching β-strands are coloured in green. Additional examples are shown in Extended Data Fig. 9b. Note that predicted coiled-coil structures are sometimes bent due to the presence of short regions of non-coiled-coil structure, which may be SFP attachment sites. **b,** Structure-based sequence alignment of predicted β-prism structures of the F7 pyocin and diverse phage CFPs. Arrows indicate the orientation of the β-strands. Residues in positions “S”, “L” and “M” are highlighted in cyan, red and yellow, respectively. Residues that can form buried salt bridges in coiled-coil structures are colored in purple. The average pairwise sequence identity among the shown regions is 32%. A sixth strand is observed for some phages and the F7 pyocin as the switched region was longer in these cases. Phages TLS and Shlf1 are predicted to have helical regions of 40 and 33 residues, respectively, inserted into their β-prism structures between the third and fourth strands. For simplicity, the sequences of these inserts have been omitted, but a one residue gap is present in the alignment at this position to denote the presence of these insertions. **c,** Cryo-EM structure of a portion of the coiled-coil region of the F7 pyocin CFP, with residues at positions S, L, and M colored as in **b**. The inset shows the salt bridges formed between Glu1039 at an M position with Arg1043 at the i+4 position. **d,** Cryo-EM structure of the β-prism of F7 pyocin CFP.

To identify conserved sequence features that might determine the fold-switching property, we overlaid the predicted phage β-prism structures onto the solved β-prism structure of the F7 pyocin. These structures aligned well with r.m.s.d. values of 2.1 to 4.8 Å over at least 100 residues (Extended Data Fig. 9c). Structure-based sequence alignment revealed a conserved pattern of SxSxxMxxLxS in each strand of the β-prism within the fold-switching region (Fig. 5b), where “<u>S</u>mall” or “S” positions display mostly small amino acids (68% Ala, 11% Ser, 7% Thr, 8% Val), “<u>L</u>eu/Ile” or “L” positions display mostly hydrophobic residues (33% Leu, 32% Ile, 25% Val), and “Middle” or “M” positions are less conserved but display a marked preference for hydrophobic or charged residues (51% Leu, Ile, or Val; 29% Lys, Arg, Asp, or Glu). The directionality of this pattern in each strand corresponds to the orientation of the strand (arrows in Fig. 5b). Residues at the “S” positions are all buried near the corners of the β-prisms, where there is no space for larger residues, but are exposed in the coiled-coil structures (Fig. 5c,d). These residues are important for fold switching, as Gln substitutions at these positions in the F7 pyocin blocked the transition from coiled-coil to β-prism, as described above (Fig. 4; Extended Data Fig. 8). Residues at the “L” positions are buried in both the β-prisms and coiled-coil structures (Fig. 5c,d), potentially stabilizing both conformations. Residues at the “M” positions are fully exposed in the β-prisms, but buried in the coiled-coil structures (Fig. 5c,d). Notably, Asp or Glu appearing at “M” positions are often accompanied by Arg spaced 4 positions before or after (Fig. 5b,c, colored purple). These oppositely charged residues can form salt bridges that stabilize the coiled-coil structures, as observed in the F7 pyocin CFP (Fig. 5c). The importance of the “M” and “S” positions in fold switching was supported by evaluating the conformational propensities of the F7 pyocin CFP after residue substitutions at these positions, based on structures predicted by AF3 (Supplementary Text 1). Substitutions of hydrophobic residues at “M” positions with charged residues shifted the propensity toward the β-prism. Additional substitutions of small amino acids at “S” positions with Gln shifted the propensity back toward the coiled-coil (Extended Data Fig. 10). Overall, these analyses identify a conserved sequence pattern that is required for the fold switching and suggest that the fold switching mechanism is also widely employed by siphophage CFPs.

## Discussion

The structures presented here provide the first atomic-level insights into the mechanism by which an F-type pyocin, a non-contractile phage tail-like bacteriocin, penetrates the bacterial cell envelope. Our cryo-EM reconstructions reveal a dramatic fold switch in the central fiber protein (CFP), where a 163-residue trimeric α-helical coiled-coil converts to a triangular β-prism upon cell surface contact. This transition remodels the tail tip, likely leading to ejection of the tape measure protein (TMP), and the opening of a channel for membrane puncture. Mutations that destabilize the β-prism while preserving the coiled-coil abolish bactericidal activity without impairing assembly, demonstrating that the fold switch is essential for function. This mechanism contrasts with contractile tails, which rely on sheath contraction for force generation^6,7,9^, and highlights fold switching as a strategy in non-contractile phage tails. It should be noted, however, that the only siphophage tails characterized in structural detail, *E. coli* phages lambda and T5^25,29^, do not display the same extensive fold-switching phenomenon, indicating alternate modes of power generation. The switch in the F7 pyocin is likely triggered by binding to lipopolysaccharide (LPS) or other cell surface receptors, toppling a metastable pre-ejection state, and releasing stored conformational energy to drive TMP ejection and membrane disruption.

AlphaFold^11^ predictions imply that coiled-coil to β-prism fold switches similar to the F7 pyocin are conserved in diverse siphophage tail homologues (Fig. 5b and Extended Data Fig. 10). Structural alignments of the F-pyocin β-prism and predicted phage prism structures revealed a conserved SxSxxMxxLxS motif (Fig. 5b). The S positions in this motif are buried in the β-prism corners, but exposed in the coiled-coil, the “L” positions are buried in both folds, while the “M” position is buried only in the coiled-coil. The existence of this motif provides clues as to how a single conserved sequence pattern can be compatible with two structurally distinct conformations. Further modeling and mutagenesis studies on such fold-switching motifs will reveal principles for designing these types of proteins for use for other biotechnological applications.

The large-scale conformation changes observed in the F7 pyocin and phage tail tips are reminiscent of viral fusion proteins, such as influenza hemagglutinin and HIV Env, which exploit large-scale conformational rearrangements to bring about membrane fusion^41,42^. For example, the conformational change of hemagglutinin includes a ∼35 residue loop region in the pre-fusion state switching its fold to a helical coiled coil in the post-fusion state^43^. In the cases of phages and eukaryotic viruses, large conformational changes appear to provide a conserved biophysical strategy for storing and releasing energy needed to breach membranes without ATP hydrolysis or dedicated motor proteins.

The fold switch described here exemplifies an important means of converting chemical potential energy into mechanical work. Unlike motor proteins or ATPases, the energy is pre-loaded during biosynthesis, enabling rapid, irreversible activation upon environmental cues. Our data indicate that this fold switching has been selectively maintained across phage evolution, and we anticipate that fold switching may be employed for energy storage in other unexplored viral and microbial systems. Our findings also reveal engineering principles for designing nanoscale, nucleic acid-free bacteriocins. By harnessing programmable fold switches, synthetic tails could be tailored for targeted antimicrobial activity against resistant pathogens. Future work could explore modulating switch energetics for controlled release in therapeutic delivery systems, advancing phage-inspired nanotechnologies for infection control and beyond.

## Methods

### Bacterial strains and plasmids

All strains, pyocins, and plasmids are listed in the Supplementary Table 4. Strains were cultured in lysogeny broth (LB) or on LB agar. Plasmids were transformed into *P. aeruginosa* using electroporation and chemically competent *E. coli* by heat shock as previously described^44^. When required, gentamycin antibiotic supplementation was used at concentrations of 50 μg/mL for *P. aeruginosa* and 30 μg/mL for *E. coli*.

### Pyocin production and purification

Overnight culture of RYC97083283 (Accession #: CP199844) carrying the F7 pyocin were inoculated into fresh LB (1% inoculum used) and grown to OD_600_ = 0.5 at 37 °C. An inducing agent, Mitomycin C, was added to a final concentration of 4 μg/mL and shaking was resumed at 30 °C until cultures lysed. To ensure complete bacterial lysis, chloroform was added to induced cultures (1**–**2 drops/mL) and allowed to shake for an additional 15 minutes at 30 °C. Cultures were spun at 10,000 × *g* for 15 minutes to pellet bacterial debris. 50 μg/mL Proteinase K, 10 μg/mL DNase I and 10 μg/mL RNase A were added to lysed cultures and incubated at room temperature for 30 minutes. Supernatants containing F7 pyocins were stored at 4 °C for further assays.

For structural analysis, the supernatant containing F7 pyocins was passed through a 0.45 μm filter and ultracentrifuged at 59,000 × *g* for 1 hour. The pellet containing F7 pyocins was resuspended in TN50 buffer (10 mM Tris-HCl, pH 7.5, 50 mM NaCl). An Amicon^®^ Ultra Centrifugal Filter with a 100 kDa cut-off was used to further concentrate F7 pyocins.

### Negative-stain electron microscopy

5 μL of pyocin lysate prepared as described above was applied to glow-discharged carbon-coated copper grids (Electron Microscopy Sciences; Carbon Film 150 Mesh, Copper) for two minutes. Samples were washed twice with double distilled water and stained with 1% uranyl acetate for 20 seconds. For cell binding samples, a single colony of PAO1 were resuspended in 50 μL of TN50 buffer. 10 μL of cell mixture was incubated with 10 μL of pyocin lysate for 30 minutes at 37 °C, and then applied to glow-discharged carbon-coated grids for staining. Grids were imaged on either a ThermoFisher Talos L120C transmission electron microscope equipped with a 4k × 4k CMOS camera or a Hitachi HT7800 TEM equipped with a Xarosa 20 Megapixel CMOS camera.

### Cryo-EM specimen preparation and data collection

To prepare cryo-EM samples, 3 µL of the purified F7 pyocin sample was applied to a glow-discharged lacey gold grid coated with a supporting ultrathin carbon film (01824G, Ted Pella). The grid was blotted using filter paper (595 filter paper, Ted Pella) for 20 s with blot force 8 at 10°C and 100% humidity, and flash-frozen in liquid ethane using an FEI Vitrobot Mark IV. Cryo-EM grids were loaded into a ThermoFisher Titan Krios electron microscope operated at 300 kV for automated data collection using SerialEM^45^. Movies of dose-fractionated frames were acquired at a nominal magnification of 81,000× with a Gatan K3 direct electron detector, corresponding to a calibrated pixel size of 0.55 Å on the specimen level in super-resolution mode. A Gatan Imaging Filter (GIF) was inserted between the electron microscope and the K3 camera and operated at zero-loss mode with the slit width of 20 eV. The defocus was set between −0.8 μm and −3.0 μm. The total dose rate on the sample was set to ∼50 electrons/Å^2^, which was fractionated into 40 frames. A total of 41,609 movies were collected in three imaging sessions with cryo-EM grids made from the same batch of sample (Extended Data Fig. 1a).

### Cryo-EM data processing

The cryo-EM data processing workflow is outlined in Extended Data Fig. 1b. 41,609 collected movies were pre-processed on-the-fly using cryoSPARC Live^46^ (v4.5.1) during data collection sessions, including steps of patch motion correction, patch CTF (contrast transfer function) estimation, and exposure curation. After discarding micrographs with ice contamination, defocus value outside the range from −0.8 to −3.0 μm, or CTF fit resolution worse than 7 Å, 36,805 good micrographs were selected and then further processed using cryoSPARC^46^. Around 500,000 particles were initially picked from 4,000 representative micrographs using template-free blob picker and screened by 2D classification. Particles from good classes with clear features for the terminator, tube, fiber, or tip (Extended Data Fig. 1c) were selected to train corresponding particle detection models in Topaz^47^. These models were used for particle picking from the entire dataset. The picked particles were extracted in boxes of 400 pixels square and 2× binned to 200 pixels square (pixel size of 2.2 Å) for multiple rounds of 2D classification.

For the fiber complex, 505,377 particles were selected after 2D classification and re-extracted in boxes of 400 pixels square without binning (pixel size of 1.1 Å). These particles were subjected to *ab-initial* reconstruction (three initial models asked) and heterogeneous refinement jobs. 359,107 particles were selected and refined to 3.1 Å resolution with C3 symmetry after homogeneous refinement, CTF refinement, reference-based motion correction, and non-uniform refinement^48^.

For the terminator complex, 201,736 particles were selected after 2D classification and re-extracted in boxes of 256 pixels square without binning (pixel size of 1.1 Å). 112,045 particles were further selected after heterogeneous refinement and then refined to 3.1 Å resolution with C6 symmetry.

For the tip complex at the pre-ejection state, 412,196 particles were selected after 2D classification and re-extracted in boxes of 450 pixels square without binning (pixel size of 1.1 Å). 112,611 particles were further selected after heterogeneous refinement and then refined to 2.9 Å resolution with C3 symmetry.

For the tip complex at the post-ejection state, 435,273 particles were selected after 2D classification and re-extracted in boxes of 256 pixels square without binning (pixel size of 1.1 Å). 141,966 particles were further selected after heterogeneous refinement and then refined to 3.0 Å resolution with C3 symmetry. To resolve the asymmetric region of CFPs, these particles were re-extracted in boxes of 400 pixels square (pixel size of 1.1 Å), refined with C3 symmetry, and expanded three times by C3 symmetry. The resulting 425,898 particles were subjected to 3D classification without alignment using a sphere mask covering one side of the tip (Extended Data Fig. 1b). After screening, 52,498 particles from the class with clear features for the goblet-shaped prism were selected and refined to 3.6 Å without applying any symmetry.

For the tube complex, 3,285,638 non-overlapping particles were selected after 2D classification and re-extracted in boxes of 256 pixels square without binning (pixel size of 1.1 Å). These particles were refined to 3.2 Å resolution with C6 symmetry and then subjected to 3D classification without alignment using a soft mask covering the two layers of tube proteins in the middle. 261,467 particles from the best class were selected and refined to 2.9 Å with C6 symmetry. To investigate the flexibility of tube, 3D variability analysis (3DVA) ^15^ was conducted with 3,285,638 tube particles (Supplementary Video 2). The tube particles were clustered into 20 groups based on the 3DVA result. 201,037 particles from the group with the most pronounced curvature were selected and refined to 3.3 Å resolution without applying any symmetry. The resulting cryo-EM map of the curved tube, along with the maps of terminator, tip, and fiber in the pre-ejection state, was used to generate the composite map of F7 pyocin shown in Fig. 1a.

The overall and local resolutions of all cryo-EM maps were estimated using cryoSPARC. The maps were auto-sharpened using cryoSPARC with negative B-factors. Data collection and processing statistics were given in Supplementary Table 1.

### Model building and refinement

All atomic models were built in Coot (version 0.9.8.7) ^49,50^. PyoF2 (tail tube protein, TTP) was built *de novo* according to the cryo-EM density map of the “straight” tube at 2.9 Å resolution. The C-terminal helix of PyoF5 (tape measure protein, TMP, residues 579–611), PyoF6 (distal tail protein, DTP), PyoF7 (tail hub protein, THP), the C-terminal domain of PyoF9 (tail hub internal protein, THIP, residues 128–200 for the pre-ejection state and 158–200 for the post-ejection state), and the N-terminal domains of PyoF10 (central fiber protein, CFP, residues 5–816 for the pre-ejection state and 25–617 for the post-ejection state) were built *de novo* against the maps of the tip complexes. The C-terminal coiled-coil region of CFP (residues 921–1083) and PyoF13 (side fiber protein, SFP) were built *de novo* against the map of the fiber complex. With cryo-EM density maps at around 3 Å resolution, we were able to clearly and confidently assign not only the backbone α-carbon positions, but also side chain identities.

After placing two layers of TTPs into the cryo-EM density map of the terminator complex, we found that the densities at its distal end did not match any known protein within the F-type pyocin gene cluster. To identify this “unknown” protein, we used cryoID^51^ and ModelAngelo^19^ to build an initial model without providing any sequence information. The protein sequence was inferred from the high-resolution side chain densities in the resulting model. This sequence was used as a query in a BLAST search of the translated genome of the F7 pyocin producing strain (RYC97083283) to identify the gene encoding this unknown protein, which we named TrP_PyoF_. The model of TrP_PyoF_ was rebuilt using the exact genome encoded protein sequence.

*De novo* modeling was not applied to the C-terminal prism domain of CFP (residues 819–1222) in the post-ejection state, as well as its linkers to the tip (residues 618–818), due to the 5–7 Å local resolution. These models were generated using AlphaFold3^27^, rigid-body fitted into the asymmetric density of the post-tip, and refined manually in Coot^49,50^.

Atomic models were refined through an iterative process combining automatic refinement in Phenix (version 1.20.1) ^52^ with default parameters and manual corrections in Coot (version 0.9.8.7) ^49,50^. Initially, individual proteins were refined against the cryo-EM maps to improve secondary structure, Ramachandran and rotamer restraints, and intramolecular clash scores. At each refinement step, models were manually inspected for quality, adjusted as needed, and re-refined iteratively until a final structure was obtained. The refined proteins were then assembled in Coot^49,50^ to generate full models of the terminator, tube, fiber, and tip complexes. All models underwent additional refinement using ISOLDE^53^ to resolve clashes and geometry outliers, followed by a final refinement in Phenix^52^. The completed models were validated in Phenix^52^ and visualized using ChimeraX^54,55^. Refinement statistics are summarized in Supplementary Table 1.

### Pyocin spotting assays

150 μL of overnight culture of PAO1 was added to 4 mL of 0.6% LB agar and top plated on a 1.5% agar plate. Pyocin lysates were tenfold serially diluted in TN50 buffer (10 mM Tris-HCl, pH 7.5, 50 mM NaCl) and spotted on the surface. Plates were incubated at 37 ^°^C overnight.

### Quantitative pyocin cell-killing assays

Overnight cultures of PAO1 were inoculated into fresh LB (1% inoculum used) and grown to OD_600_ = 0.5 at 37 °C. 1 mL of culture was pelleted at 10,000 × *g* for 2 minutes and resuspended in TN50 buffer. Cultures were serially diluted in TN50 buffer. 100 μL of diluted culture was treated with 10 μL of F-type pyocin lysate and incubated at 37 ^°^C for 1 hour. The entire reaction was plated onto LB agar plates and incubated overnight at 37 ^°^C to allow for colony growth.

### Time-lapse fluorescence microscopy

Overnight culture of PAO1 was inoculated into fresh LB (1% inoculum used) and grown to OD600 = 0.5 at 37 °C before pelleting and resuspending in TN50 buffer. Cells were supplied with a final concentration of 30 μM propidium iodide (BioShop, PPI777.10) and treated with pyocin lysate. Cells were imaged on a Zeiss Axio Observer spinning disc microscope equipped with a fluorescent light source (HXP 120V, Zeiss) and a 63x/1.4NA oil objective (Zeiss) every 3 minutes. The samples were maintained at 37 °C during imagining in an incubation chamber (Zeiss). Acquisitions were made using the Zen 2 Blue software (Zeiss). Downstream analysis was performed with ImageJ v1.54f^56^.

### DNA cloning and manipulation

In-frame deletion mutants of the F7 pyocin TrP and SFPs were generated using a deletion construct in PexG2 (Accession #: KM887143) as previously described^57^. 550 nucleotide homology arms, amplified using PCR from the strain RYC97083283, were used to facilitate homologous recombination to result in in-frame, internal deletions of TrP amino acids 49–118 and SFPs amino acids 101–180. PexG2 constructs were transformed into *E. coli* strain SM10-λpir for mating into RYC97083283. Merodiploid cells were streaked onto LB agar plates (no salt) with 15% (w/v) sucrose. All mutants were confirmed via PCR and DNA sequencing performed by the Centre for Applied Genomics (The Hospital for Sick Children, Toronto, Canada).

Amino acid substitutions of F7 pyocin CFP residues were similarly introduced into the strain RYC97083283 by homologous recombination. First, wild-type CFP with 400 nucleotide homology arms (CFP residues L772–R1214) were amplified by PCR from RYC97083283 and cloned into PexG2 using restriction enzymes. Using this construct as a template, single residue mutations were created by site-directed mutagenesis and recombined into RYC97083283 as described above. F7_CFP-4Q_ and F7_CFP-10Q_ were similarly created using homologous recombination. CFP genes harbouring the 4 and 10 Gln substitutions were synthesized by Twist Bioscience (South San Francisco, CA, USA) and ligated into PexG2. All cloning and mutagenesis were confirmed via PCR and DNA sequencing performed by the Centre for Applied Genomics (The Hospital for Sick Children, Toronto, Canada).

### Quantification of pyocin tail length

Negative-stain electron micrographs were used to count the layer number of hexameric TTP rings constituting the pyocin tail. The length of the tail tube (excluding tail tip, terminator and fibers) was calculated by multiplying the layer number of TPP rings with the average length of each ring. 82 F7_WT_ and 86 F7_ΔTrP_ pyocin particles were counted.

### Bioinformatics

Phage TrPs, PyoF11 and PyoF12 homologs, and phage CFPs bearing the coiled-coil region were identified using BLAST^39^ searches. Multiple sequence alignments were generated in Jalview (version 2) ^58^ using MUSCLE^59^ with default settings. Pairwise alignment percent identities were generated using Jalview^58^. Protein structure comparisons were produced using DALI^60^. Conservation of the TrP of the F7 pyocin was confirmed by BLAST^39^ searches against *Pae* strain sequences in the Integrated Microbial Genomes and Microbiomes (IMG/M) database^61^.

To assemble an initial set of phage CFP sequences with potential fold-switching regions, we searched an in-house annotated phage genome database^62^ with the C-terminal region (residue 816 to the end) of the F7 pyocin CFP. From a database of 120 siphophages infecting Gram-negative bacteria, we identified 45 related CFPs (e-value < 1e-6). Redundant sequences (pairwise sequence identity > 80%) were eliminated to bring the number further characterized to 24.

### Cryo-ET sample preparation

*P. aeruginosa* strain PAO1 was grown on an LB agar plate at 37 °C overnight. A single isolated colony was picked and resuspended in 50 μL of filter sterilized phosphate buffer saline (PBS). Then, 5 μL of resuspended bacteria was spotted onto at least five different spots on a new LB agar plate. The plate was incubated at 37°C for at least 12 hours. *P. aeruginosa* at the edge of the spots was harvested and resuspended in TN50 buffer (10 mM Tris-HCl, pH 7.5, 50 mM NaCl) to a final OD_600_ of 1.6. In a fresh microcentrifuge tube, 4 μL of pyocins was added into 16 μL of the bacterial sample, followed by incubation at 37 °C for 10 min. Similarly, to examine the later stage of pyocin infection and capture F7 pyocins in the post-ejection state, four-fold diluted pyocin sample was used to infect the strain, followed by incubation at 37 °C for one hour. 16 μL of BSA coated gold tracer solution with 10 nm particle size (Aurion) was added to the infected *P. aeruginosa* samples. For cryo-ET sample preparation, 5 μL of the mixture was deposited onto freshly glow-discharged cryo-EM grids (Quantifoil, Cu), followed by back blotting using filter paper (Whatman^TM^) for 4 seconds. The grids were then immediately plunged into liquid ethane and propane mixture using a manual gravity plunger, as described previously^63^.

### Cryo-ET data collection

Frozen-hydrated specimens of *P. aeruginosa* with pyocin were imaged below -180°C using Titan Krios G2 300kV transmission electron microscope (ThermoFisher) equipped with a field emission gun, a K3 direct detection camera (Gatan), and a GIF BioQuantum imaging Filter (Gatan). SerialEM software^45^ was used for data collection. Tilt series were acquired at a magnification of 42,000×, corresponding to a physical pixel size of 2.148 Å. The dose-symmetric scheme in FastTomo script^64^ was used to tilt the specimen stage of the microscope from −48° to +48° in increments of 3°. The total electron dose was approximately 65 e^-^/Å^2^ distributed across 33 images in each tilt series.

### Reconstruction of tomograms

Image drift induced by the electron beam during data acquisition was corrected by MotionCor2^65^. Image stacks were generated using IMOD^66^. Approximately 10 fiducial beads in each tilt series were tracked using IMOD^67^ for tilt series alignment. For tilt series with fewer fiducial beads, patch tracking in IMOD was used for alignment. Gctf^68^ and the ctfphaseflip functions in IMOD^69^ were used for defocus determination and CTF correction, respectively, for all images in the aligned tilt series. For particle picking, the aligned tilt series were 6× binned using the binvol function in IMOD^66^. Then, 6× binned tomograms with simultaneous iterative reconstruction technique (SIRT) were reconstructed by Tomo3D^70^. Pyocin particles in the pre- and post-ejection states were manually picked from the tomograms, as described previously^71–73^. 4,683 particles were picked for the initial stage of pyocin binding to bacterial membrane, and 124 particles were picked for the late stage of binding.

### Subtomogram-averaged structure of the pyocin bound to the *P. aeruginosa* cell surface

For particle picking, the aligned tilt series were 6× binned using the binvol function in IMOD^66^. Then, 6× binned tomograms with simultaneous iterative reconstruction technique (SIRT) were reconstructed by Tomo3D^70^. Pyocin particles in the pre- and post-ejection states were manually picked from the tomograms, as described previously^71–73^. 4,683 particles were picked for the initial stage of pyocin binding to bacterial membrane, and 124 particles were picked for the late stage of binding. For subtomogram averaging, 6× binned tomograms with weighted back projection (WBP) were reconstructed by Tomo3D^70^. Then, protomo software package^71,72^ was used for subtomogram alignment and averaging. Based on the aligned position of pyocin in the 6× binned tomograms, unbinned subtomograms were extracted, and 2× and 4× binned subtomograms were subsequently generated using the binvol function in IMOD^66^. To improve structural resolution, the averaged structures were refined using the 4× binned subtomograms. The 3D classification was performed to remove bad particles and to determine C3 symmetrical features in class-averaged structures of pyocin in the pre-ejection state. For the structure in the pre-ejection state, 2× binned subtomograms were used for the refinement process. Then, C3 rotational symmetry was applied to the averaged structure to improve structural resolution. For the structure in the post-ejection state, 2× binned subtomograms were used for the refinement process. Then, rotational symmetry was applied to the averaged structure to improve signal-to-noise ratio.

### Molecular modeling of the *in-situ* pyocin structure bound to the *P. aeruginosa* cell surface

The 3D rendering images were created using ChimeraX^54,55^, and structures of pyocin and the outer membrane were segmented out from the maps. The high-resolution structures of pyocin determined by single-particle cryo-EM were then fitted into the *in-situ* structures.

### Structural prediction using AlphaFold

Initial structural predictions on phage CFP sequences were performed using AlphaFold3^27^ using default settings on the web server (https://alphafoldserver.com/). Predictions of F7 pyocin CFP mutants were performed on the web server using 10 different non-random starting seed structures.

ColabFold1.5.5^74^, an efficient-yet-accurate implementation of AlphaFold2^75^, was run on the NIH HPC Biowulf cluster (https://hpc.nih.gov/apps/colabfold.html) to generate multiple sequence alignments (MMseqs2-based routine) and all predictions. Initial predictions of alternative conformations were run using CF-random^11^ on the full sequence of wild-type F-pyocin with the AlphaFold2 multimer model (v3). At full MSA sampling depth (--max-seq=512, --max-extra-seq=5120, notation 512:5120 following), only the prism structure was predicted. A mixture of coiled-coil and prism structures was predicted sampling 64:128, both with moderate-to-high average per-residue Local Distance Difference Test (pLDDT) confidences ≥ 70.

ColabFold1.5.5^74^ was used to generate multiple sequence alignments (MMseqs2-based routine) and predictions for the sequence-diverse 24 CF phages on the NIH HPC Biowulf cluster (https://hpc.nih.gov/apps/colabfold.html). Structures of only the fold-switching and neighboring regions–comprising residues ∼800–1200–were predicted. The parameters for each run were as follows: “--num-seeds 5 --num-models 5 --model-type alphafold2_multimer_v3”. The predictions were generated using the full/deep Multiple Sequence Alignments (MSA), and were also sampled at 512:5120, 64:128, and 32:64, where X:Y denotes the --max-seq and --max-extra-seq flags, respectively.

To detect fold-switching in the sequence diverse set of 24 phage CFPs, the following steps were taken: 1) The predictions were compared with the experimentally determined folds of the prism and coiled-coiled helix using MMalign (https://zhanggroup.org/MM-align/) and overall, Root Mean Square Deviations (RMSD) were calculated. 2) MM-scores > 0.5 were filtered to create a list of potential hits that matched both folds. Low RMSD values (< 20 Å) were detected. 3) Visual inspection of structures was used the MM-scores were close to the threshold of 0.5. The scripts used for all analyses were written in Python, Bash, and PyMOL Version 3.0 (Schrodinger, https://pymol.org/) was used to visualize the proteins.

### Quantification and statistical analysis

Data were presented as means (s.d.). calculated from three biological replicates. The number of F7 pyocin tail tube units, and number of assembled F7 pyocins were counted manually from negative-stain EM images, and statistics were determined using GraphPad Prism version 8.4.2 (GraphPad Software, Boston, MA, USA, https://www.graphpad.com). All the details can be found in the figure legends and method details.

## Supplementary Text 1

To test the importance of the “M” and “S” positions in fold switching, we used AF3^27^ to predict the structures of the F7 pyocin CFP sequence with amino acid substitutions at some of these positions. AF3^27^ consistently predicted the wild-type F7 pyocin CFP switch region as a coiled-coil (Extended Data Fig. 10a). Since the “M” positions in our structural alignment are predicted to be important for stabilizing the coiled-coil but not the β-prism, we expected that destabilizing substitutions at these positions in the pyocin CFP sequence would switch the structure to a β-prism. Consistent with this prediction, a mutant in which three hydrophobic residues at “M” positions were substituted with charged residues (V1007R/ L1022D/I1054E, F7_CFP-3M_) was predicted by AF3^27^ as all β-prism in 7 out of 10 structures generated using 10 different non-randomly chosen seed sequences (Extended Data Fig. 10b). Having created a mutant form of the F7 pyocin CFP that was consistently predicted as a β-prism, we aimed to determine whether substitutions at “S” positions, which are predicted to be crucial only for maintaining the β-prism structure, would discourage β-prism formation within the context the F7_CFP-3M_ mutant. For this purpose, we tested the Gln substitutions described above that were designed to block fold switching and experimentally found to abrogate biological activity. These substitutions are all in the “S” positions. Strikingly, introducing the 4 Gln substitutions into the β-prism-forming mutant background (F7_CFP-3M/4Q_) generated structures that all had decreased levels of β-prism and increased levels of coiled-coil structure, though the amount varied between structures (Extended Data Fig. 10c). Models predicted for the mutant with only 2 Gln substitutions at “S” positions (F7_CFP-3M/2Q_) also displayed a marked shift towards coiled-coil containing structures (Extended Data Fig. 10d), but to a lesser extent than the F7_CFP-3M/4Q_ mutant. It is interesting to note that even though AF3^27^ effectively predicted changes in fold-switching propensity resulting from amino acid substitutions, it was not nearly as effective as CF-random in detecting fold-switching for the wild-type phage sequences. Overall, this modeling of F7 pyocin CFP mutants supports the importance of the conserved sequence pattern seen in the phage switch regions and demonstrates that the mutants we tested experimentally have a reduced propensity to adopt the β-prism structure.

## Acknowledgements

This study was supported by grants from the US National Institutes of Health (R01GM071940 and DE025567 to Z.H.Z., GM124378 and R01AI087946 to J.L.) and a grant from the Canadian Institutes of Health Research to A.R.D. (FDN-15427). A.R.D. is the Canada Research Chair in Bacteriophage-Based Technologies (CRC-2024-00209). L.L.P. was supported by the Intramural Research Program of the National Institutes of Health (NIH, LM202011). We acknowledge the use of resources at the Electron Imaging Center for Nanosystems [supported by UCLA and by instrumentation grants from the NIH (1S10OD018111) and NSF (DBI-1338135 and DMR-1548924)] and the Yale CryoEM resource (funded in part by the NIH grant 1S10OD023603-01A1). The contributions of the NIH author(s) are considered Works of the United States Government. The findings and conclusions presented in this paper are those of the author(s) and do not necessarily reflect the views of the NIH or the U.S. Department of Health and Human Services.

## Author contributions

A.R.D, Z.H.Z and J.L. conceived the study. L.L.P. supervised the structural predictions. Y.H., A.S.C.L., X.C., S.T., R.K. and D.C. performed experiments. Y.H., A.S.C.L., X.C., A.R.D., Z.H.Z. wrote the paper. All authors edited and approved the paper.

## Competing interests

We declare there is no competing interest.

## Materials availability

Materials generated by the authors in this study will be distributed upon request.

## Data and code availability

The cryo-EM density maps were deposited in the Electron Microscopy Data Bank (EMDB) with accession code: tail fiber in the pre-ejection state (EMD-72915), tail terminator (EMD-72916), tail tube (EMD-72917), curved tail tube (EMD-72918), tail tip in the pre-ejection state (EMD-72919), tail tip in the post-ejection state (EMD-72920), asymmetric reconstruction of tail tip in the post-ejection state (EMD-72921), *in situ* structure of tail tip in the pre-ejection state (EMD-72402), and *in situ* structure of tail tip in the post-ejection state (EMD-72401).

The corresponding models were deposited in the Protein Data Bank (PDB) with tail fiber in the pre-ejection state (9YG8), tail terminator (9YG9), tail tube (9YGA), tail tip in the pre-ejection state (9YGB), tail tip in the post-ejection state (9YGC), and asymmetric structure of tail tip in the post-ejection state (9YGD).

This paper does not report any original code.

Any additional information required to reanalyze the data reported in this paper is available from the lead contact upon request.

---

**Extended Data Fig. 1.**
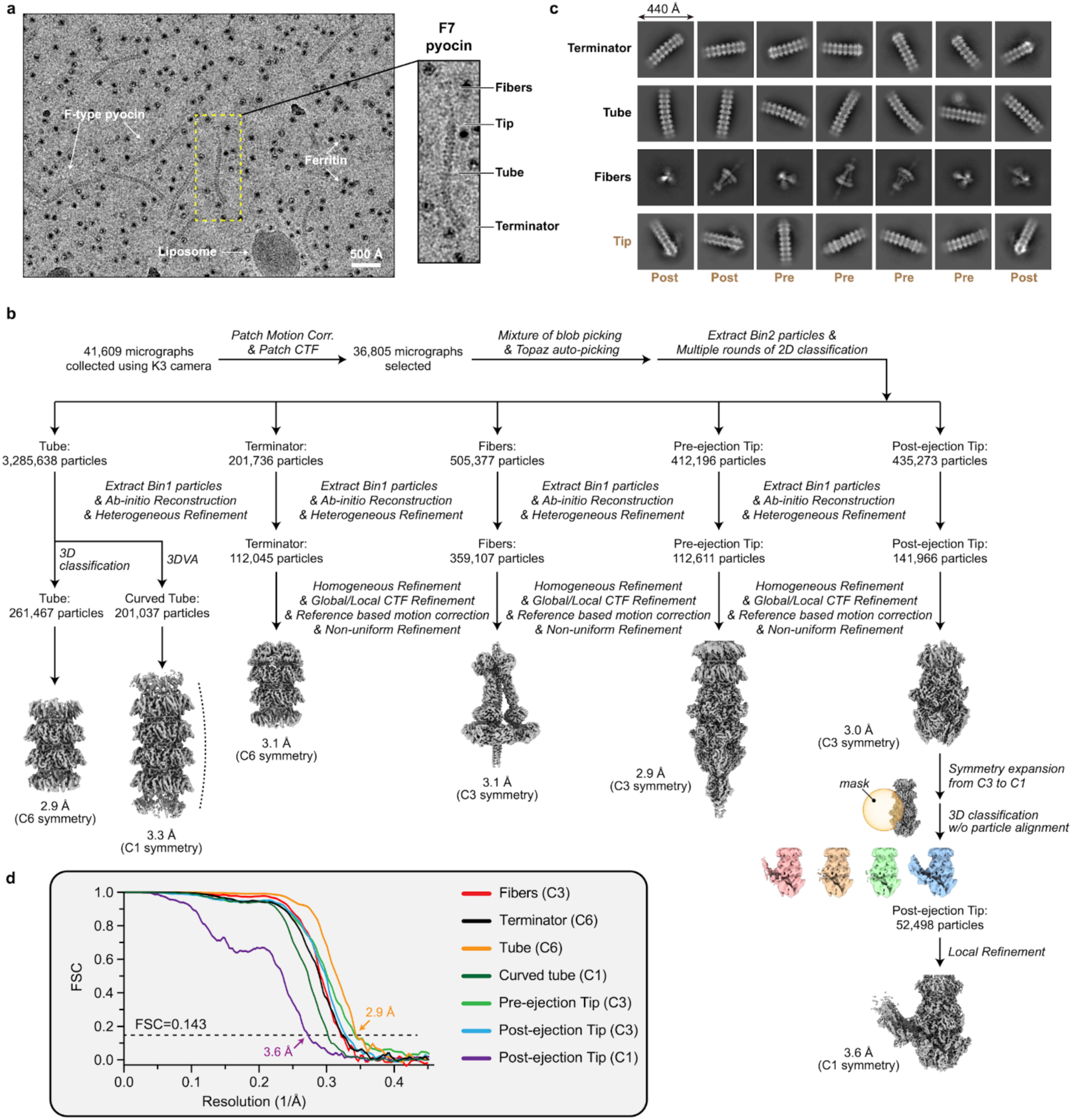
| Cryo-EM structure determination of F7 pyocin. **a,** A representative cryo-EM image of the endogenously purified F7 pyocin sample. Pyocin particles, along with particles of ferritin and liposome, are labeled. The zoom-in view of one pyocin particle is shown on the right, with its structural components labeled. **b,** Cryo-EM data processing workflow (detailed in Methods). **c,** Representative 2D class averages of the terminator, tube, fibers, and tip (in both pre- and post-ejection states) regions of F7 pyocin. **d,** Plot of the Fourier shell correlation (FSC) as a function of the spatial frequency demonstrating the resolutions of final reconstructions.

**Extended Data Fig. 2.**
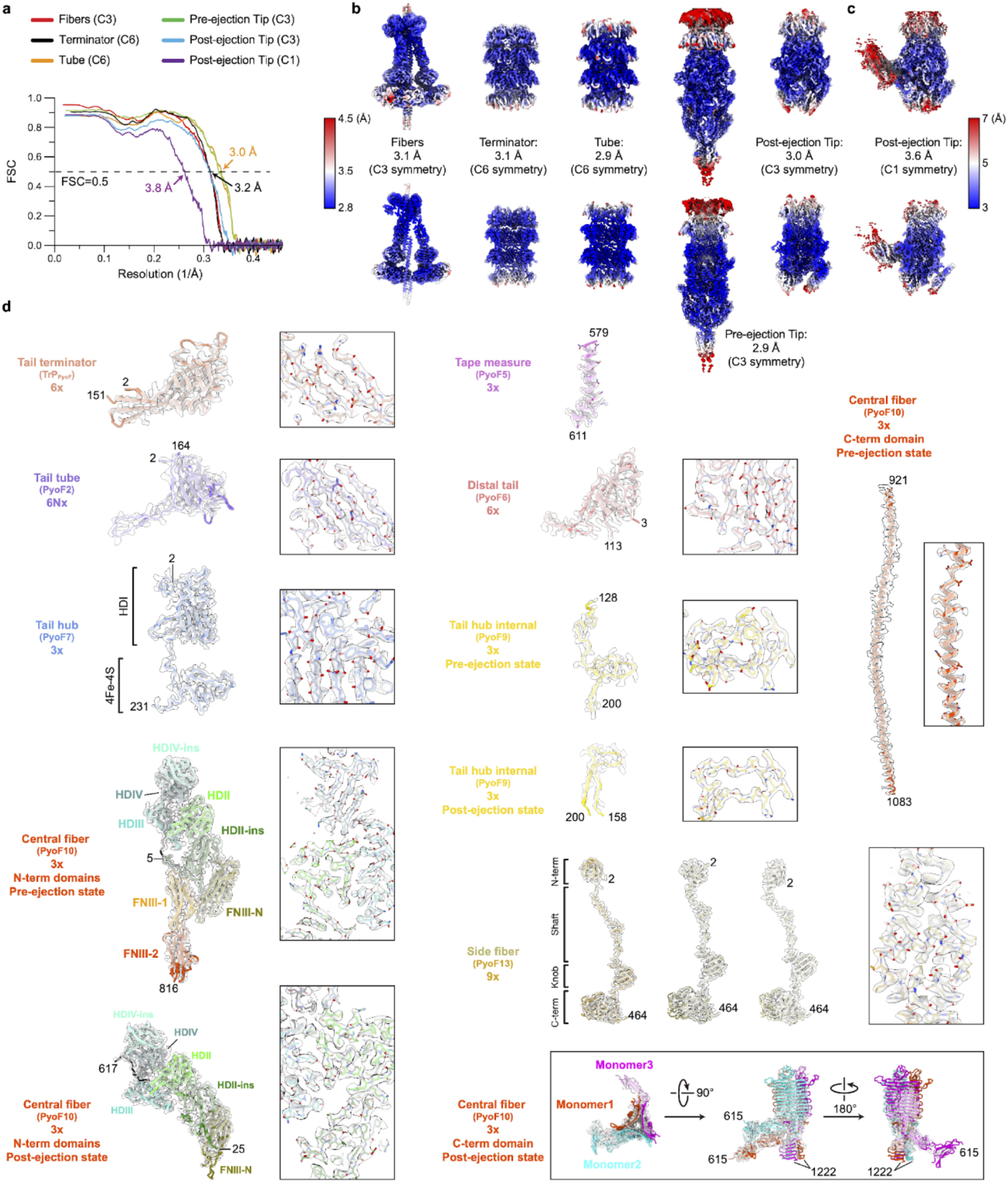
| Evaluation of cryo-EM density maps and models of F7 pyocin. **a,** FSC coefficients as a function of spatial frequency between models and corresponding cryo-EM density maps. **b–c,** Local resolution evaluation of cryo-EM density maps. Note that maps in (**b**) and (**c**) are colored using different color codes. **d,** Cryo-EM densities encasing the corresponding atomic models of individual proteins. Numbers denote chain termini. Regions of the cryo-EM density map (semi-transparent densities) superimposed with atomic models (sticks) are shown in boxes, demonstrating the agreement between the observed and modelled amino acid side chains.

**Extended Data Fig. 3.**
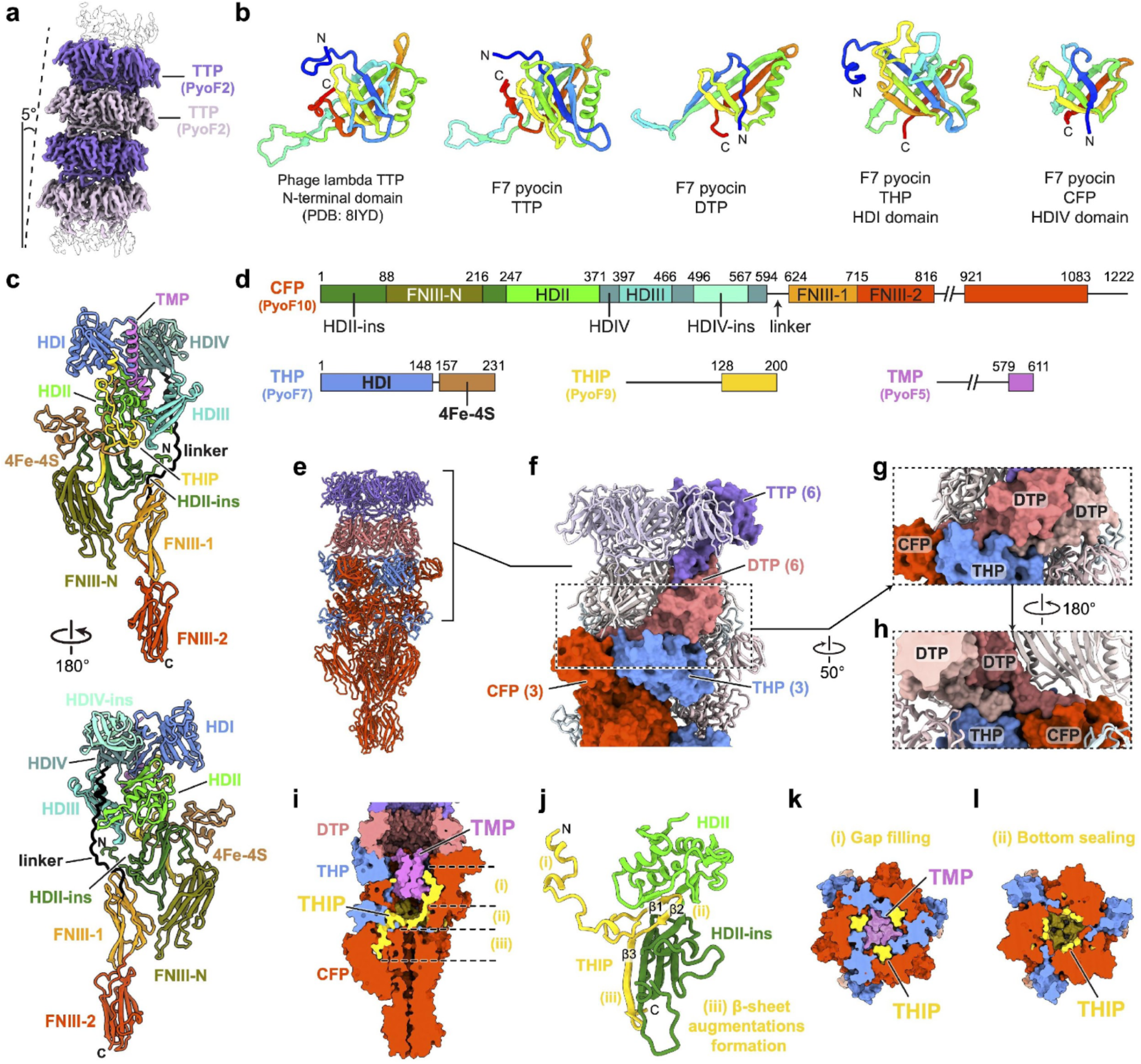
| Molecular organization of the tail tube and tail tip. **a,** Cryo-EM density map of the curved tail tube. **b,** Rainbow-colored ribbon diagrams of the N-terminal domains of phage λ TTP^13^, F7 pyocin TTP, F7 pyocin DTP, the HDI domain of F7 pyocin THP, and the HDIV domain of F7 pyocin CFP, highlighting the conserved tail tube fold. **c-d,** Ribbon diagram (**c**) and linear schematic (**d**) of the domain organization of CFP, THP, THIP, and TMP. **e–f,** Molecular organization of the tail tip of F7 pyocin. The DTP ring functions as a symmetry adapter, connecting the C6 symmetrical tail tube to the C3 symmetrical tail tip. **g–h,** Detailed interactions between DTP and THP-CFP viewed from the surface side (**g**) and lumen side (**h**). **i–l,** Location and interactions of THIP in the lumen of tail tip. The three regions of THIP (i, ii, and iii) perform distinct functions.

**Extended Data Fig. 4.**
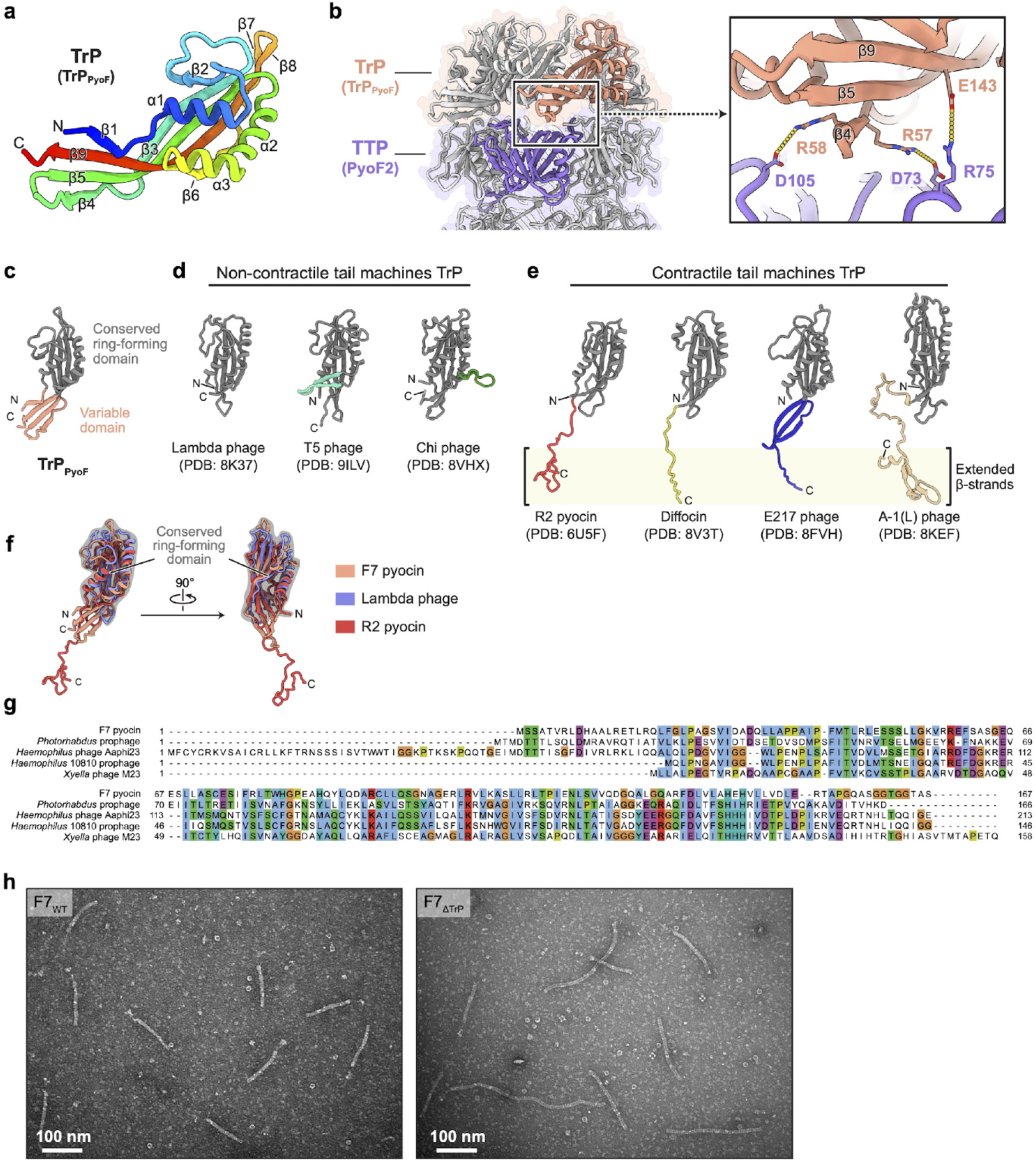
| Structural and sequence comparisons of tail terminators from various phage tail-like bacteriocins and phages. **a,** Rainbow-colored ribbon diagram of TrP from F7 pyocin. **b,** Interactions between the TrP ring and the TTP ring. **c–e,** Structural comparison of TrPs from F7 pyocin (**c**), phage lambda^12^, phage T5^79^, phage Chi^14^, R2 pyocin^6^, diffocin^7^, phage E217^80^, and phage A-1 (L) ^81^. The conserved ring-forming domains are colored in grey, and the variable domains are colored in different colors. The extended β-strands in **e** facilitate interaction with the sheath proteins of contractile phage tails and contractile phage tail-like bacteriocins. **f,** Structural alignment of TrPs from F7 pyocin, phage lambda^12^, and R2 pyocin^6^. **g,** Multiple sequence alignment of TrPs of the F7 pyocin and diverse phages. Phages were identified through BLAST^39^ searches using the sequence of *PA0910* as a query. **h,** Representative negative-stain EM images of F7_WT_ and F7_ΔTrP_ mutant particles.

**Extended Data Fig. 5.**
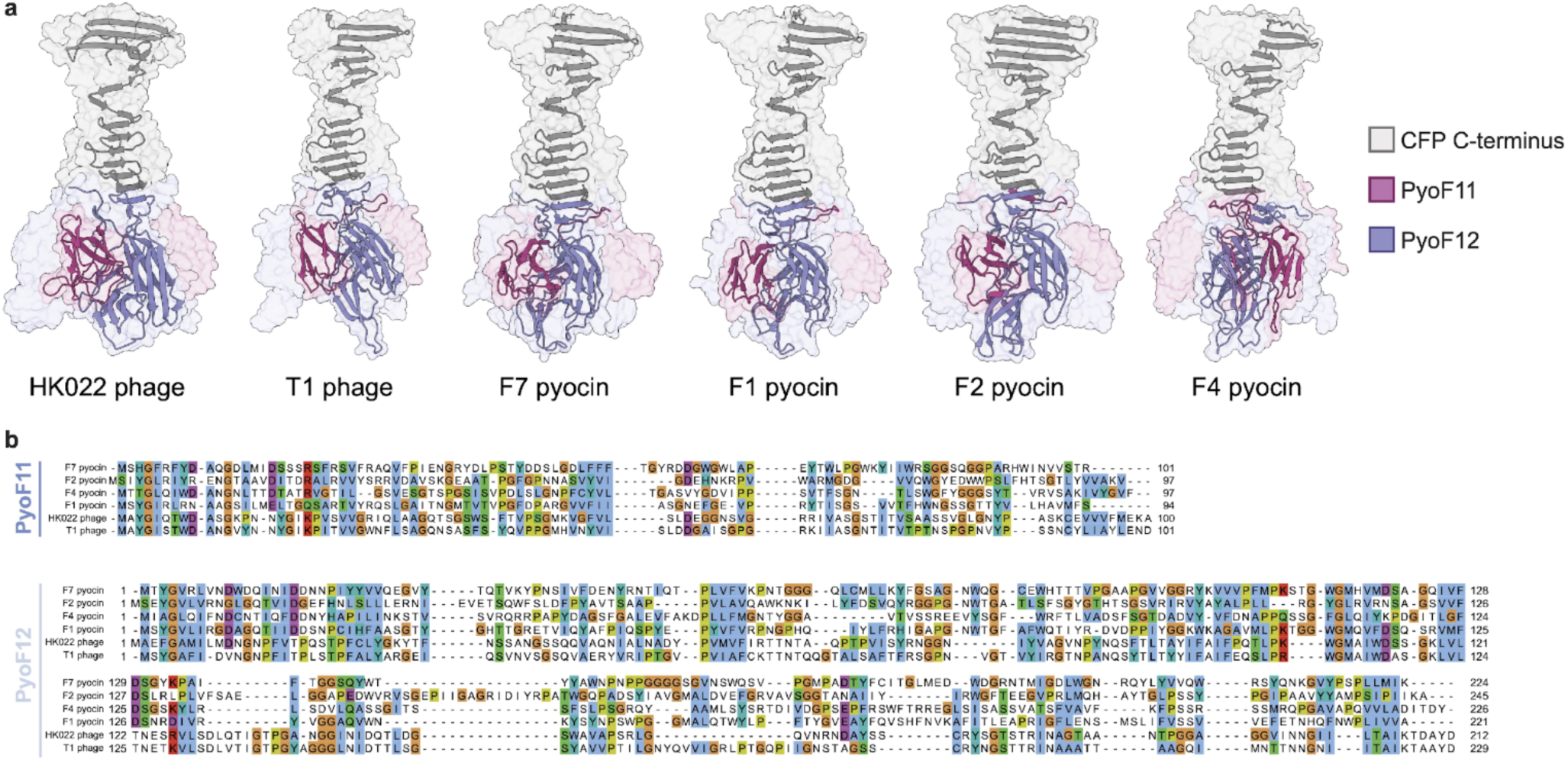
| PyoF11 and PyoF12 homologs can be identified in F-type pyocin groups and diverse phages. **a,** AlphaFold3 (AF3) ^27^-predicted structures of the CFP:PyoF11:PyoF12 complexes for various F-type pyocin groups and phages. **b,** Multiple sequence alignments of PyoF11 and PyoF12 from different F-type pyocin groups and homologs from diverse phages.

**Extended Data Fig. 6.**
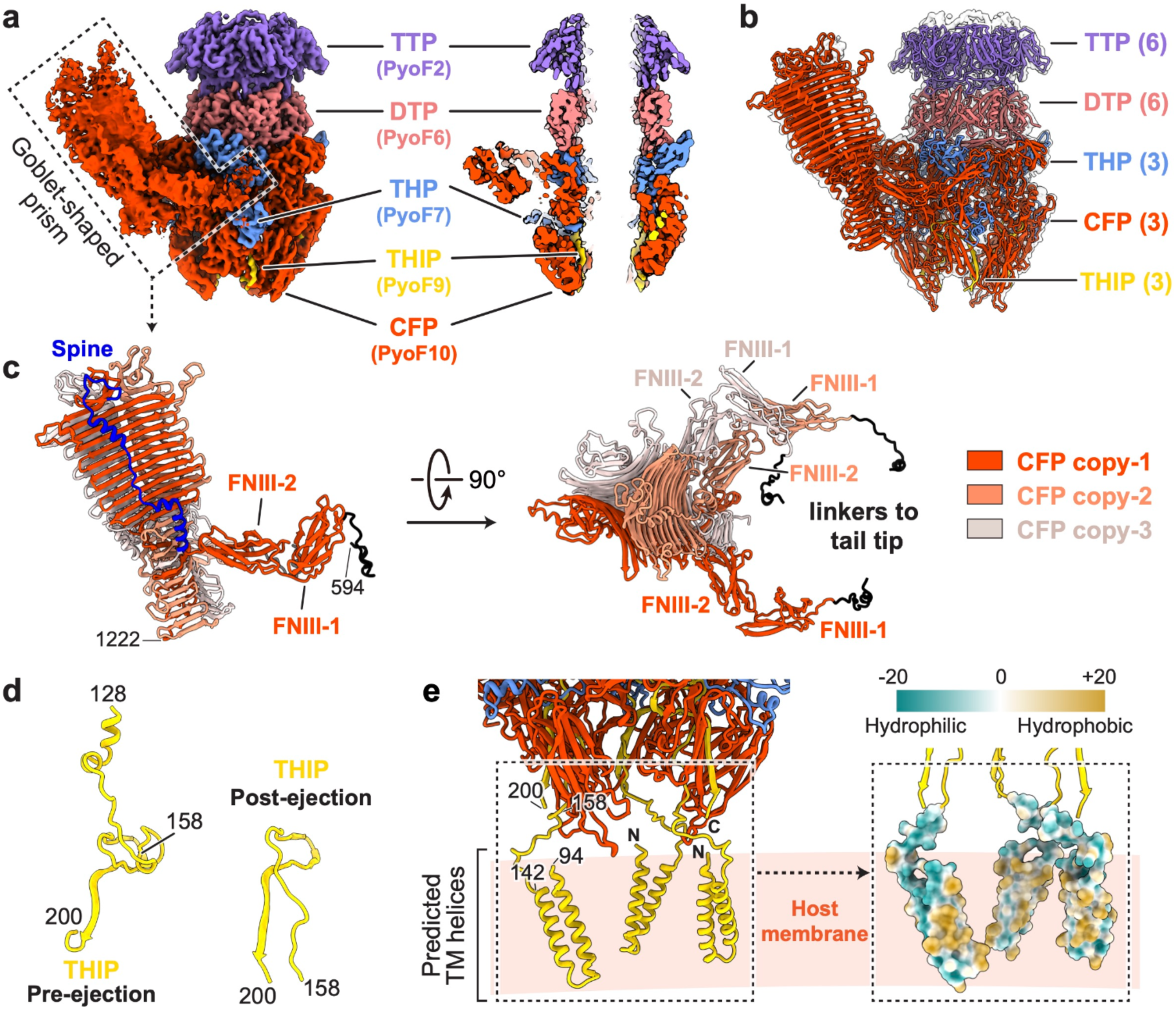
| Structure of the tip and fiber complex in the post-ejection state. **a,** Cryo-EM density map of the tail tip and fiber complex in the post-ejection state, with the sectional view shown on the right. **b,** Atomic models of the tip and fiber complex in the post-ejection state. **c,** Side and top views of the β-prism of CFP trimer, as framed by dotted lines in (**a**). The three CFP subunits are colored in red, coral, and light pink, respectively. **d,** Ribbon diagrams of THIP in the pre-ejection and post-ejection states. **e,** The potential transmembrane (TM) helices in THIP. Residues 94–142 are predicted as two helices by AF3^27^ and as the TM region by TMHMM 2.0^28^. The hydrophobic surface of the TM helices is shown on the right.

**Extended Data Fig. 7.**
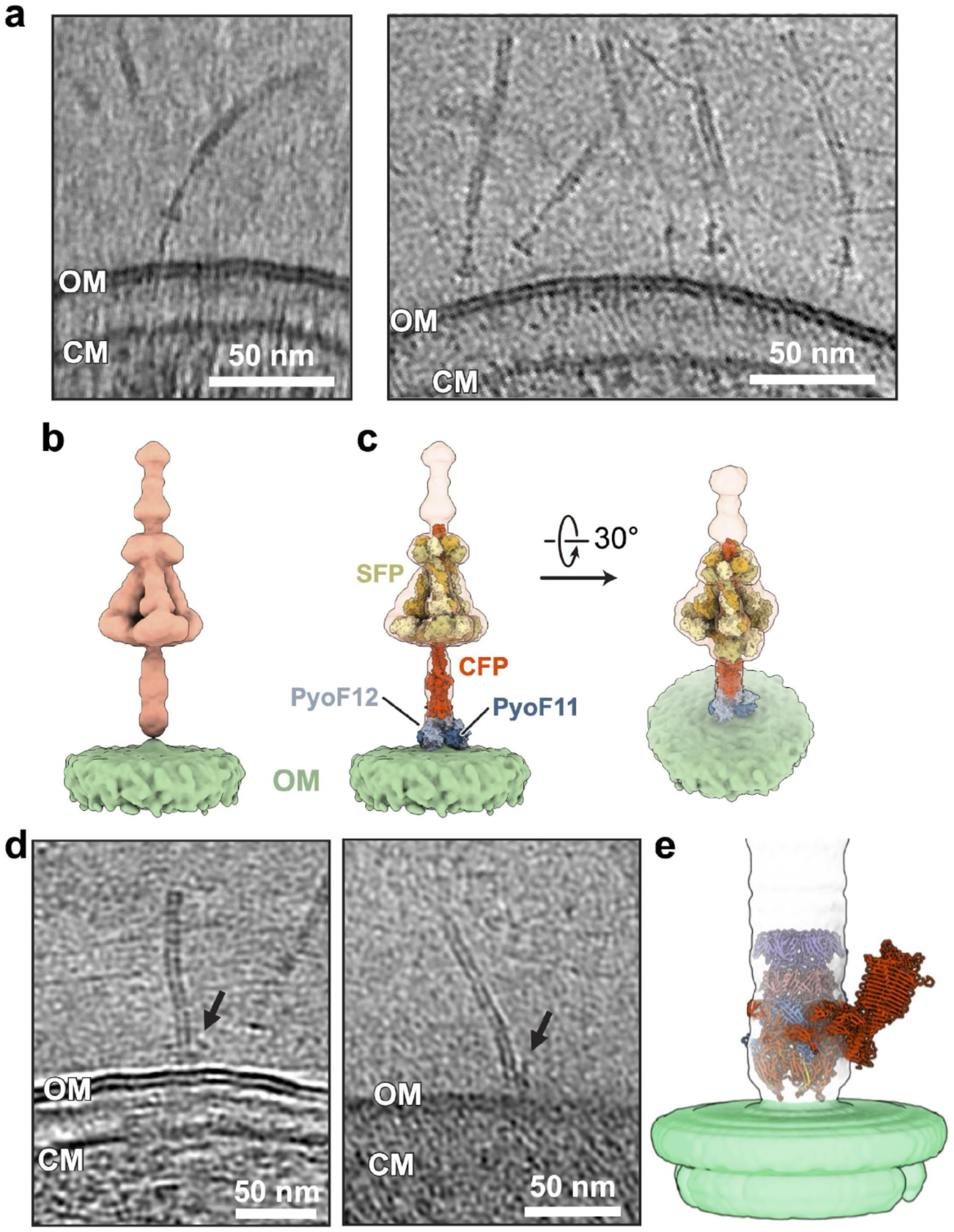
| Reconstructed tomograms and subtomogram-averaged density maps of F7 pyocin on the surface of *P. aeruginosa*. **a,** Slice images of cryo-ET reconstructions of F7 pyocin in the pre-ejection state. **b–c,** Subtomogram-averaged density map of F7 pyocin in the pre-ejection state, shown alone (**b**) and overlaid with the model of tail fiber (**c**). **d,** Slice images of cryo-ET reconstructions showing F7 pyocin in the post-ejection state. The density corresponding to the CFP β-prism is indicated by an arrow. OM, outer membrane. CM, cell membrane. **e,** Subtomogram-averaged density map of F7 pyocin in the post-ejection state fitted with the model of tip complex. The β-prism is not seen because refinement was performed with C3 symmetry. The density of outer membrane is colored in light green.

**Extended Data Fig. 8.**
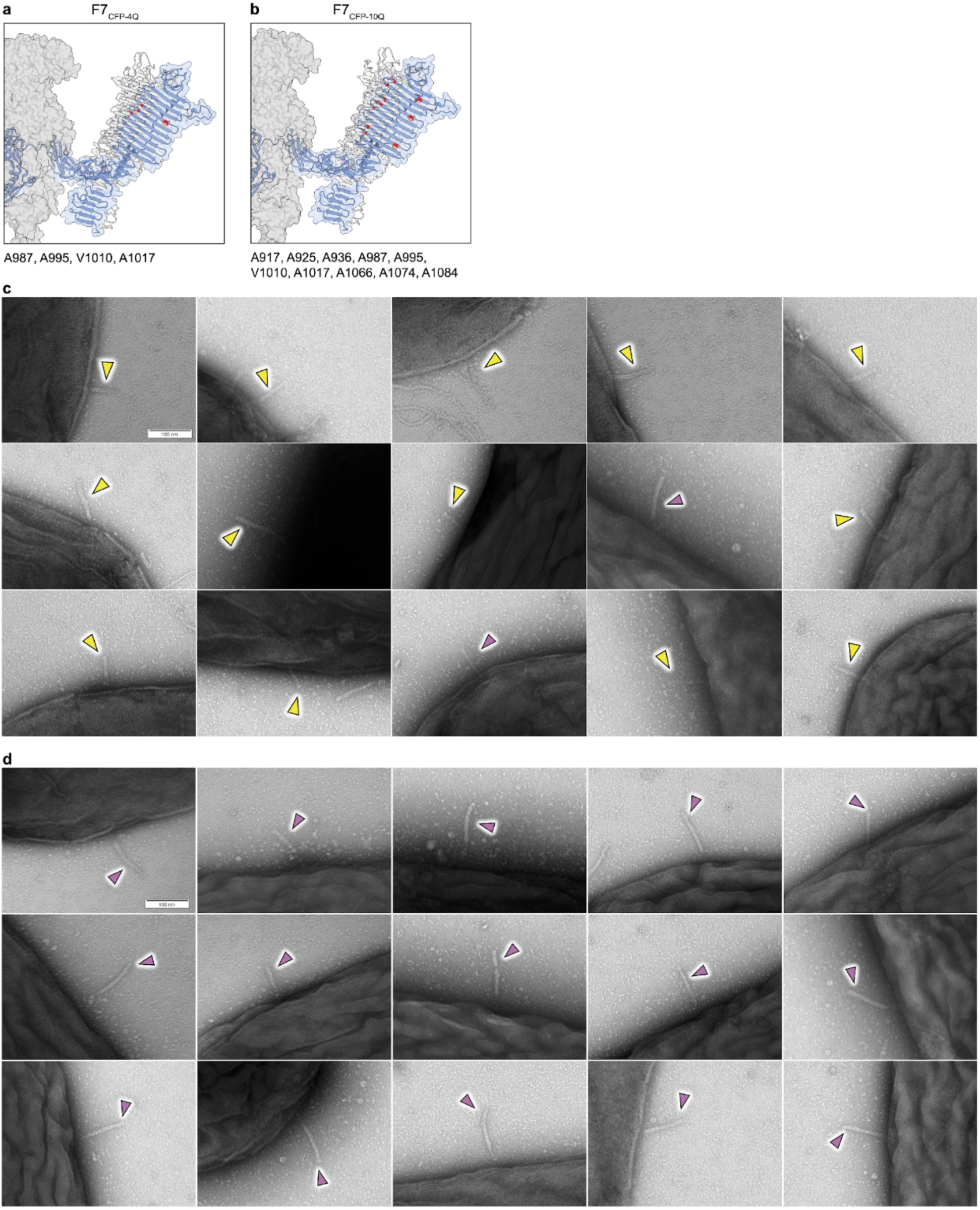
| Amino acid substitutions in the coiled-coil region to inhibit fold-switching. **a–b,** The locations of the mutated residues in the constructs F7_CFP-4Q_ (**a**) and F7_CFP-10Q_ (**b**) in the β-prism structure. **c–d,** Additional electron micrographs of F7_WT_ (**c**) and F7_CFP-2Q_ mutant (**d**) interacting with strain PAO1, related to Fig. 4i–j. Yellow and purple arrows indicate pyocins lacking TMP and those containing TMP in the tube lumen, respectively.

**Extended Data Fig. 9.**
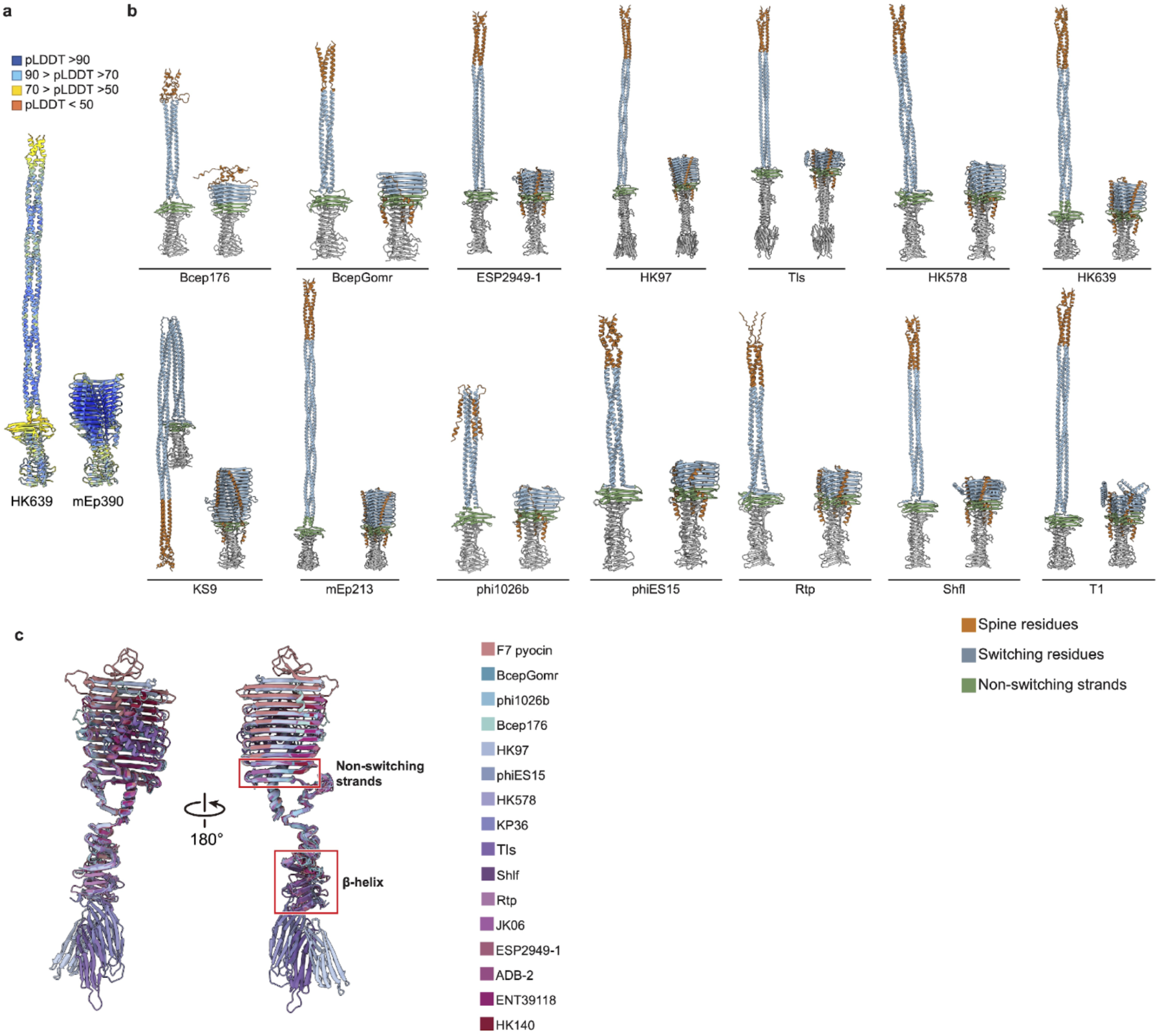
| Predicted fold-switched structures of diverse phage CFPs, showing coiled-coil to β-prism transitions. **a,** AF3^27^ predictions of HK639 and mEp390 phage CFP C-terminal regions, which are 80% identical in sequence. **b,** Fold-switched structures of phage CFPs predicted by CF-random^11^, related to Fig. 5a. The non-switching strands are colored in green. Residues that fold switch from coiled-coil to β-prism are colored in blue. Residues forming the spine in the β-prism are colored in dark orange. Note that predicted coiled-coil structures are sometimes bent due to the presence of short regions of non-coiled-coil structure, which may be SFP attachment sites. **c,** Structural comparison of β-prisms of phage CFPs predicted by CF-random.

**Extended Data Fig. 10.**
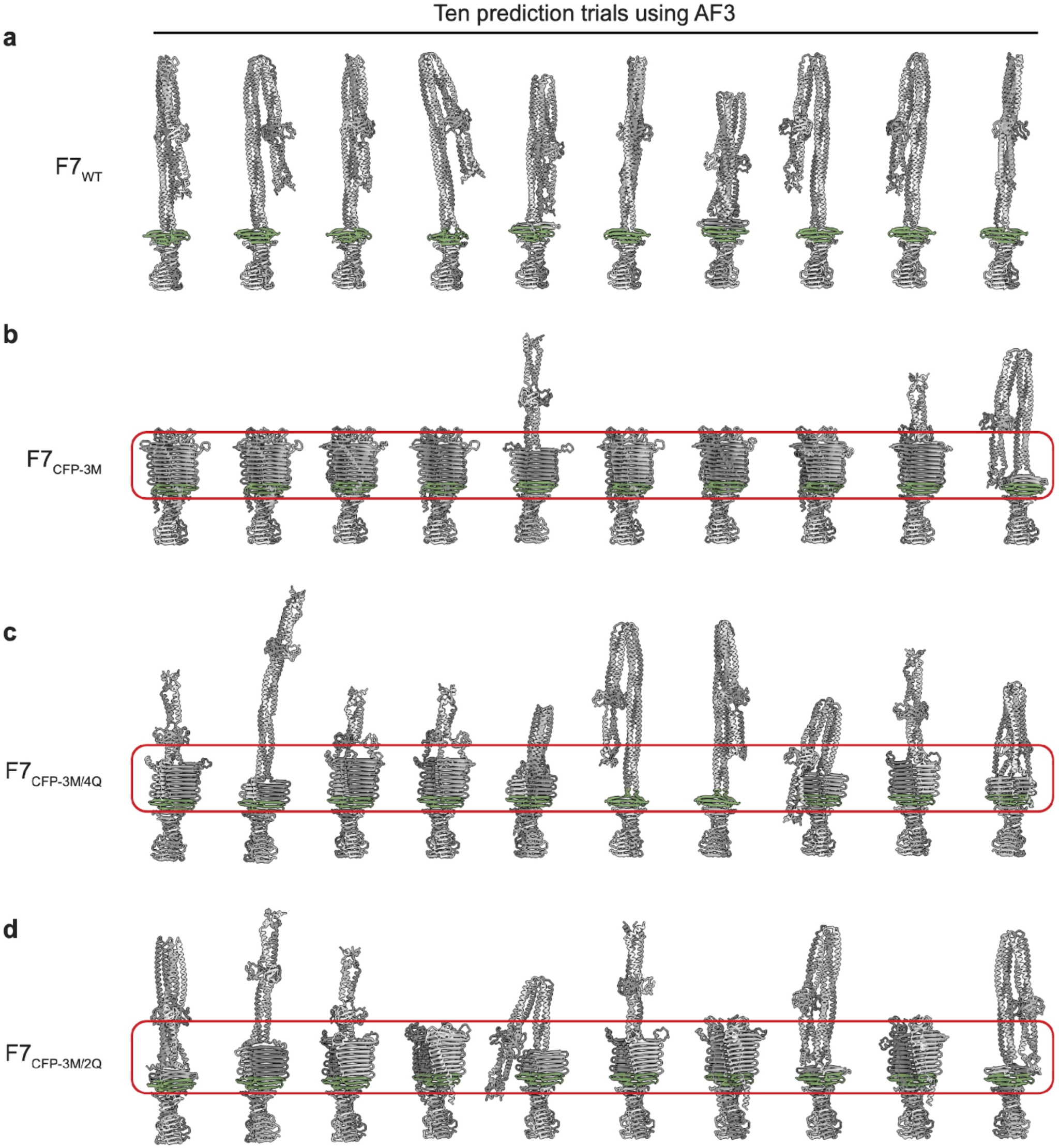
| AF3 predictions of F7 pyocin CFP switch region mutants. **a–d,** Residues 818–1222 of the F7_WT_, F7_CFP-3M_, F7_CFP-3M/4Q_, and F7_CFP-3M/2Q_ pyocins were predicted by AF3^27^, respectively. For the WT and each mutant, AF3 was run 10 times using a different specified seed each time. In this way, the 10 distinct structures for each case are shown here. The three non-switching prism strands that are present in both coiled-coil and β-prism structures are shown in green. For comparison between structures, the red rounded rectangles denote the extent of the complete β-prism structure.

**Supplementary Fig. 1.**
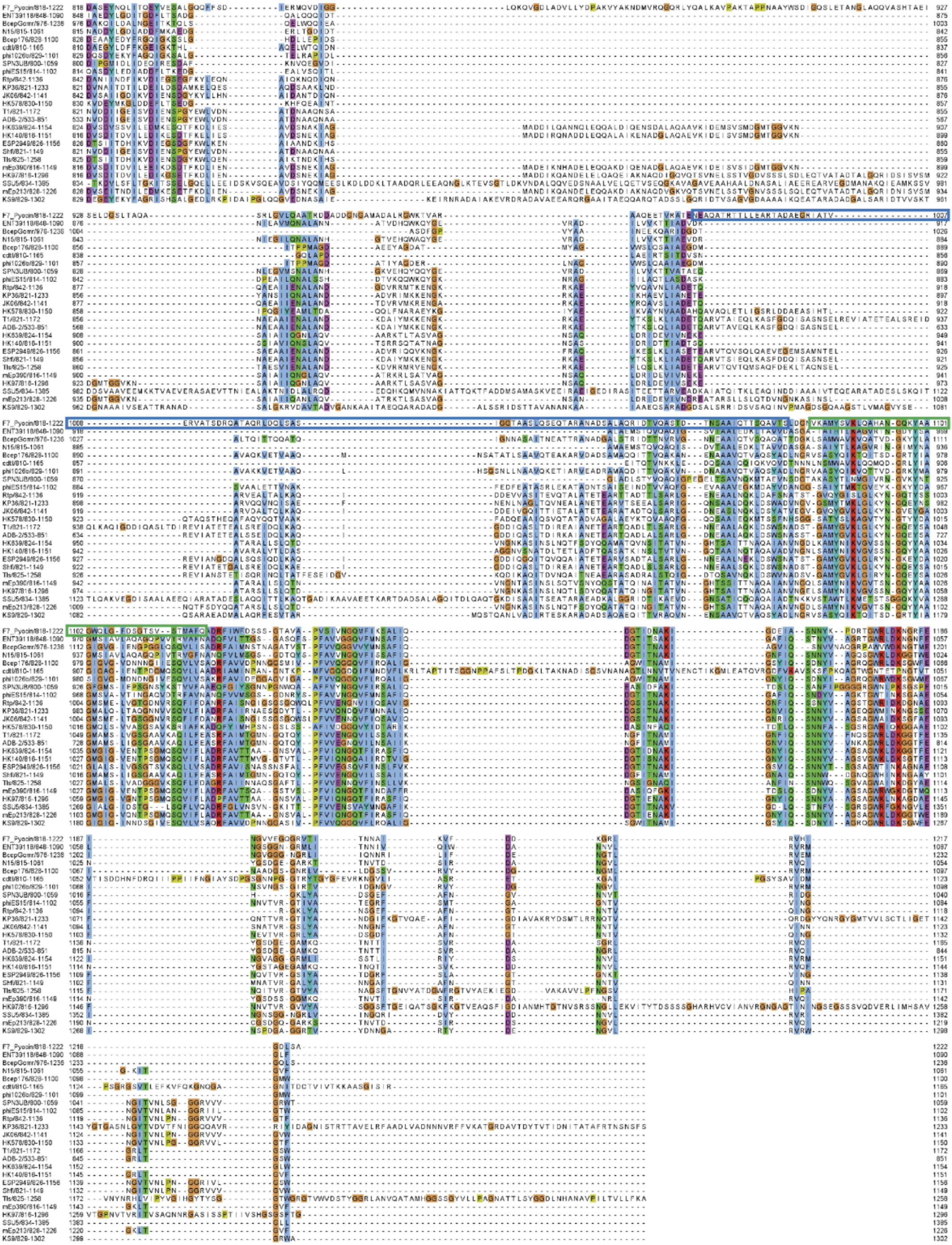
| Sequence of 24 phage CFPs predicted to switch from coiled-coil to β-prism. The regions shown begin approximately 20 residues N-terminal to the start of the fold-switching region and extend to the end of the proteins. Fold-switching and non–fold-switching regions are highlighted with blue and green frames, respectively.

**Supplementary Table 1.**
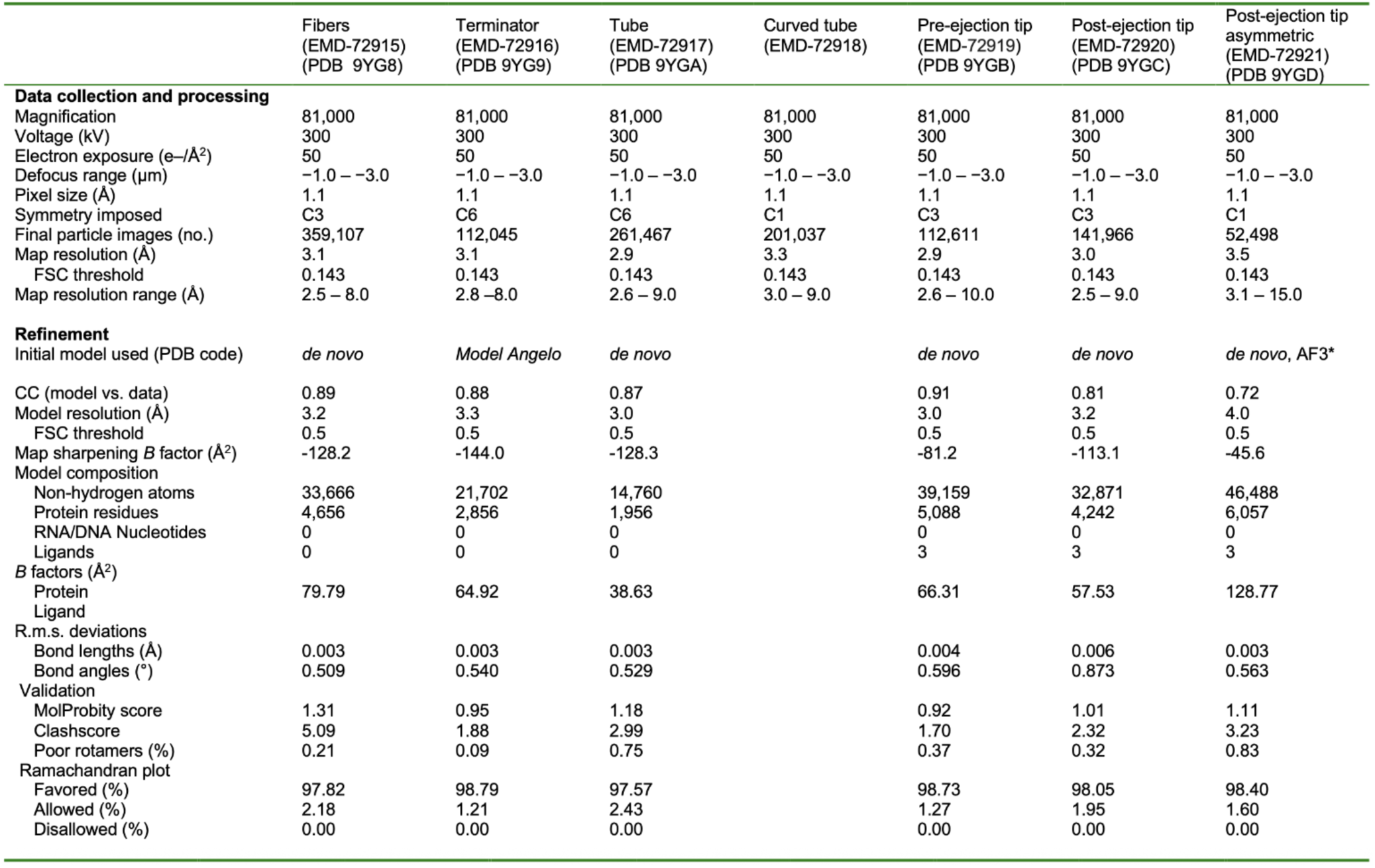
Cryo-EM data collection, refinement and validation statistics.

**Supplementary Table 2.** Cryo-ET data collection, refinement and map generation.

| Microscope parameters | Pre-ejection state | Post-ejection state |
| --- | --- | --- |
| Microscope | TFS Krios | TFS Krios |
| Voltage (kV) | 300 | 300 |
| Magnification | 42,000x | 42,000x |
| Physical pixel size (Å) | 2.148 | 2.148 |
| Total electron dose / tilt series (e<sup>-</sup>/Å) | 65 | 65 |
| Tilt angles | +48° ~ - 48° | +48° ~ - 48° |
| Tilt increments | 3° | 3° |
| Target defocus (μm) | -4.8 | -4.8 |
| Detector | BioQuantum K3 (Gatan) | BioQuantum K3 (Gatan) |
| Energy Filter slits (e<sup>-</sup>) | 20 | 20 |
| Tomography / STA | Pre-ejection state | Post-ejection state |
| Tomogram used | 624 | 355 |
| Initial particle numbers selected | 4654 | 122 |
| Final particle number used to reconstruct the map | 3324 | 94 |
| Symmetry applied | Rotational symmetry | Rotational symmetry |
| Map resolution (Å) | 19.0 |  |
| FSC threshold | FSC 0.5 cut-off | FSC 0.5 cut-off |
| **EMDB ID** | 72402 | 72401 |

**Supplementary Table 3.** Coiled-coil predictions for the fold-switching regions of phages.

| Predicted by CF-random |  |  |  |  |  |
| --- | --- | --- | --- | --- | --- |
| Pyocin/ phage | Coiled-coil residues predicted by Coconat | Spine residues | Non-switching residues* | Switching residues* | Central fiber protein accession number |
| F7 | 839-860, 908-951, 965-1084 | 818-862 | 1088-1120 | 903-1087 | WHT11187.1 |
| KS9 | 850-1060, 1077-1163 | 830-902 | 1166-1199 | 830-902 | YP_003090195.1 |
| HK97 | 834-1039 | 816-876 | 1042-1078 | 879-1041 | NP_037717.1 |
| mEP213 | 846-1013, 1023-1084 | 828-887 | 1090-1122 | 890-1089 | YP_007112361.1 |
| HK140 | 834-1007 | 816-871 | 958-1046 | 877-957 | YP_007111724.1 |
| mEP390 | 834-1008 | 816-871 | 981-1046 | 877-980 | YP_007112436.1 |
| N15 | 833-907 | 816-853 | No helix predicted |  | NP_046916.1 |
| HK639 | 842-1016 | 824-880 | 1022-1054 | 885-1021 | YP_004934053.1 |
| HK578 | 854-986 | 830-870 | 998-1035 | 871-997 | YP_007112629.1 |
| phiES15 | 835-938 | 814-852 | 940-987 | 856-939 | YP_006590053.1 |
| cdtI | 840-890 | 810-842 | 907-933 | 834-906 | YP_001272531.1 |
| TLS | 845-997 | 825-870 | 997-1046 | 875-996 | YP_001285540.1 |
| ESP2949 | 846-992 | 826-872 | 993-1040 | 877-992 | YP_007005423.1 |
| Shlf1 | 839-986 | 821-867 | 987-1035 | 871-986 | YP_004414866.1 |
| Rtp | 860-975 | 842-887 | 972-1023 | 891-971 | YP_398987.1 |
| JK06 | 860-973 | 842-887 | 972-1023 | 893-971 | YP_277442.1 |
| KP36 | 839-953 | 821-867 | 953-1002 | 870-952 | YP_007173568.1 |
| Bcep176 | 874-962 | 829-863 | 866-954 | 959-998 | YP_355397.1 |
| phi1026b | 871-963 | 829-866 | 964-999 | 867-963 | NP_945050.1 |
| BcepGomr | 1007-1095 | 976-1009 | 1086-1139 | 1010-1085 | YP_001210242.1 |
| ENT39118 | 866-940 | 848-866 | No helix predicted |  | YP_007238120.1 |
| SPN3UB | 806-881 | 801-838 | No helix predicted |  | YP_007011007.1 |
| T1 | 842-943, 962-972, 984-1019 | 822-868 | 1017-1068 | 869-1016 | YP_003912.1 |
| ADB-2 | 555-699 | 534-583 | 708-747 | 584-704 | YP_007112723.1 |
| SSU5 | 867-1063, 1068-1251 | N/A | No prism predicted |  | YP_006906652.1 |

\* Switching and non-switching residues were determined by overlaying predicted coiled-coil structures with predicted β-prism structures. Residues that were predicted to be a $\beta$ -strand in both the coiled-coil and β-prism structures were designated as "non-switching". Non-switching residues only include the region annotated in Extended Data Fig. 9 and do not include residues in the β-helix.

**Supplementary Table 4.** Strains, pyocins, and plasmids used in this study.

| Bacterial strain | Details | Source |
| --- | --- | --- |
| ***Pseudomonas aeruginosa*** |  |  |
| PAO1 | Wild-type | Lab collection |
| RYC97083283 (PaeA3) | Wild-type | Lab collection |
| PaeA3 (ΔTrP) | Terminator mutant, in-frame deletion | This study |
| PaeA3 (ΔSFP) | Side fiber protein mutant, in-frame deletion | Lab collection |
| PaeA3 (F7<sub>CFP-2Q</sub>) | Residue substitution (A987Q/A995Q) | This study |
| PaeA3 (F7<sub>CFP-2QC</sub>) | Residue substitution (A998Q/A1011Q) | This study |
| PaeA3 (F7<sub>CFP-4Q</sub>) | Residue substitution (A987Q/A995Q/V1010Q/A1017Q) | This study |
| PaeA3 (F7<sub>CFP-10Q</sub>) | Residue substitution (A917Q/A925Q/A936Q/A987Q/A995Q/V1010Q/A1066Q/A1074Q/A1084Q) | This study |
| ***Escherichia coli*** |  |  |
| DH5α |  | Lab collection |
| SM10 λpir | For conjugation of PexG2 into *P. aeruginosa* | Lab collection |
| **Pyocins** |  |  |
| F7 WT | Wild-type | Lab collection |
| F7<sub>CFP-2Q</sub> | F7 pyocin with residue substitutions in the central fiber protein-encoding sequence (A987Q/A995Q) | This study |
| F7<sub>CFP-2QC</sub> | F7 pyocin with residue substitutions in the central fiber protein-encoding sequence (A995Q/A1011Q) | This study |
| F7<sub>CFP-4Q</sub> | F7 pyocin with residue substitutions in the central fiber protein-encoding sequence (A987Q/A995Q/V1010Q/A1017Q) | This study |
| F7<sub>CFP-10Q</sub> | F7 pyocin with residue substitutions in the central fiber protein-encoding sequence (A917Q/A925Q/A936Q/A987Q/A995Q/V1010Q/A1066Q/A1074Q/A1084Q) | This study |
| F7ΔSFP | F7 pyocin with in-frame deletion in the side fiber protein-encoding gene | Lab collection |
| F7ΔTrP | F7 pyocin with in-frame deletion in the terminator-encoding gene | This study |
| **Plasmids** |  |  |
| PexG2 | *E. coli*- *P. aeruginosa* conjugation vector | Hmelo<sup>70</sup> |
| F7<sub>TrP</sub> | F7 pyocin terminator with homology arms to facilitate recombination cloned into the MCS PexG2 using EcoRI and HindIII | This study |
| F7ΔTrP | In-frame deletion of F7 pyocin terminator created with site-directed mutagenesis using the F7<sub>TrP</sub>-PexG2 plasmid described above | This study |
| F7<sub>CFP</sub> | F7 pyocin central fiber with homology arms to facilitate recombination cloned into the MCS of PexG2 using EcoRI and HindIII | This study |
| F7<sub>CFP-2Q</sub> | F7 pyocin central fiber with residue substitutions created with site-directed mutagenesis using the F7<sub>CFP</sub>-PexG2 plasmid | This study |
| F7<sub>CFP-2QC</sub> | F7 pyocin central fiber with residue substitutions created with site-directed mutagenesis using the F7<sub>CFP</sub>-PexG2 plasmid | This study |
| F7<sub>CFP-4Q</sub> | F7 pyocin central fiber with residue substitutions created with site-directed mutagenesis using the F7<sub>CFP</sub>-PexG2 plasmid | This study |
| F7<sub>CFP-10Q</sub> | F7 pyocin central fiber with residue substitutions created with site-directed mutagenesis using the F7<sub>CFP</sub>-PexG2 plasmid | This study |

**Supplementary Table 5.**
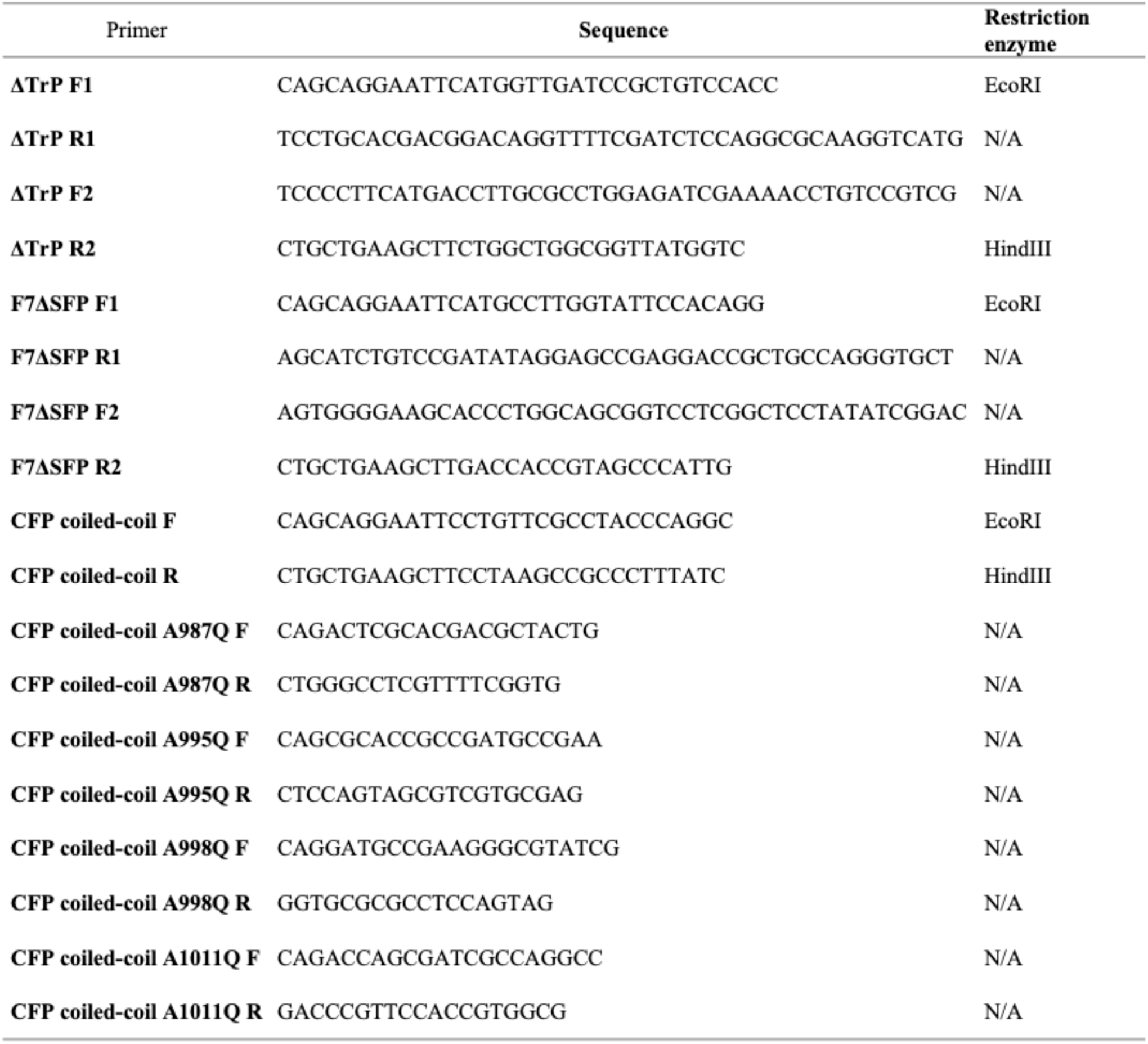
Oligonucleotides used in this study.

**Supplementary Table 6.** Key reagents and resources.

| Reagent | Source | Reference number |
| --- | --- | --- |
| Tris | Bioshop | Cat# TRS003 |
| Sodium chloride | Bioshop | Cat# SOD004 |
| BioTryptone | BioBasic | Cat# TG217 |
| Yeast extract | Bioshop | Cat# YEX555 |
| Agar | Bioshop | Cat# AGR001.500 |
| Gentamycin | Bioshop | Cat# GTA202.5 |
| Mitomycin C | Cell Signaling Technologies | Cat# 51854S |
| Proteinase K | Biobasic | Cat# PB0451 |
| DNase I | Sigma-Aldrich | Cat# 10104159001 |
| RNase A | Sigma-Aldrich | Cat# R5500 |
| EcoRI | New England Biolabs | Cat# R3101S |
| HindIII | New England Biolabs | Cat# R3104S |
| Phusion™ High-Fidelity DNA Polymerase | ThermoFisher | Cat# F530S |
| T4 Polynucleotide Kinase | New England Biolabs | Cat# M0201S |
| T4 DNA Ligase | New England Biolabs | Cat# M0202S |
| T4 DNA Ligase Buffer | New England Biolabs | Cat# B0202A |
| DpnI | ThermoFisher | Cat# ER1705 |
| Uranyl acetate | Electron Microscopy Sciences | Cat# 22400 |
| GenepHIow™ Gel/PCR Kit | FroggaBio Scientific Solutions | Cat# DFH300 |
| EZ-10 Spin Column Plasmid DNA Minipreps Kit | BioBasic | Cat# BS614 |
| Amicon® Ultra Centrifugal Filter, 100 kDa MWCO | Millipore | Cat# UFC510024 |
| 0.45 µM filter | FroggaBio Scientific | Cat# SF0.45PES |
| Carbon-coated TEM grids | Electron Microscopy Sciences | Cat# CF150-Cu |
| Quick-Seal® Round-Top Polypropylene Tube | Beckman Coulter | Cat# 342413 |
| Lacey Carbon grid with continuous ultrathin carbon | TED PELLA | Cat# 01824G |
| Blotting paper, Grade 595 | TED PELLA | Cat# 47000-100 |

**Supplementary Table 7.**
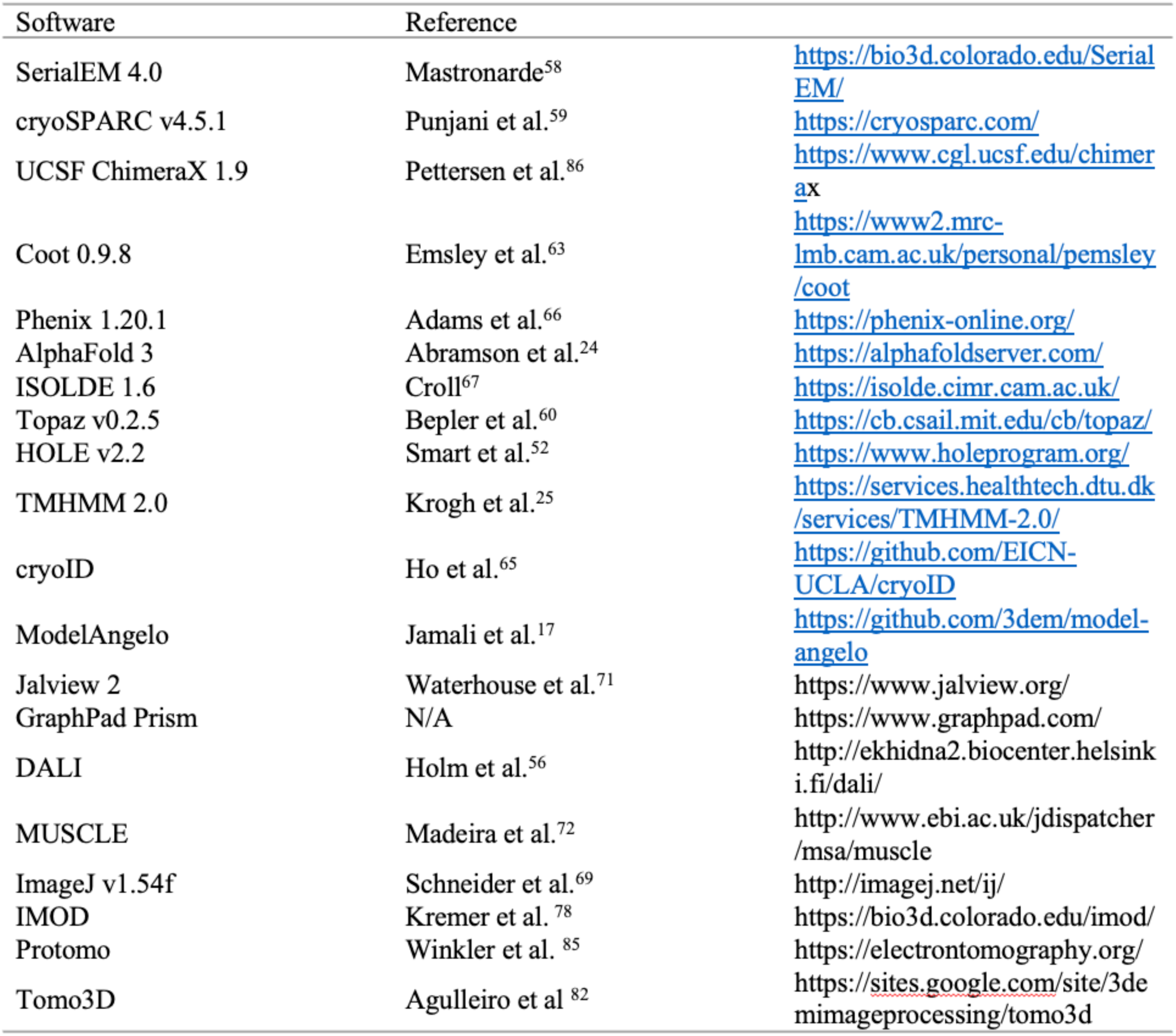
Software and algorithms.

**Supplementary Video 1 | Overall structure of F7 pyocin.**

**Supplementary Video 2 | 3D variability analysis of the tail tube of F7 pyocin.**

**Supplementary Video 3 | Structural details of the tail tip and tail fiber of F7 pyocin.**

**Supplementary Video 4 | Conformational change of the tail tip from the pre-ejection state to the post-ejection state.**

**Supplementary Video 5 | Conformational change of the tail fiber from the pre-ejection state to the post-ejection state.**

## Notes

### Competing Interest Statement

The authors have declared no competing interest.

